# Tissue-derived Activin signaling restrains regeneration after genotoxic stress in *Schmidtea mediterranea*

**DOI:** 10.64898/2025.12.29.696926

**Authors:** Haleigh Brownlee, Amit Dubey, Nirurita Mahadev, Zachary Castles, Andrea Rauschmayer, Hannah Ashraf, Blair W. Benham-Pyle

## Abstract

Regenerative capacity depends on stem cells that replace missing tissues and signals that coordinate their activity. Following damage, particularly damage that targets stem cells, regeneration often fails. Yet, whether regeneration fails after stress because surviving stem cells lack regenerative potential or because tissue signals suppress their deployment remains unclear. Here, we show that sublethal genotoxic stress reversibly inhibits whole-body regeneration in the planarian *Schmidtea mediterranea*. Following irradiation, surviving stem cells proliferate, expand, and generate lineage-biased progenitors, but fail to mount normal amputation-induced proliferative responses. Regeneration remains impaired after amputation despite stem cell recovery, but re-injury after the recovery period restores regeneration. Single-cell transcriptomic analysis revealed altered activation of transient regeneration-activated cell states (TRACS) in differentiated tissues, accompanied by increased activin ligand expression and reduced *follistatin* expression. Remarkably, depletion of *activin-2* restored amputation-induced stem cell proliferation and regeneration of missing tissues despite prior irradiation and substantial stem cell depletion. *Activin-2* depletion also improved regeneration following exposure to the chemotherapy agent cisplatin. These findings identify Activin-dependent signaling as a reversible restraint on regeneration after genotoxic stress. More broadly, they demonstrate that regeneration failure after stem cell depletion can reflect a restrictive signaling environment rather than an insufficient supply of stem cells with regenerative potential.

**In Brief:** Brownlee et al. uncover an activin/TGF-beta dependent inhibition of regenerative capacity following genotoxic stress in the highly regenerative flatworm *Schmidtea mediterranea*. While…

**Highlights:**

- Low-doses of genotoxic stress inhibit regenerative capacity for 3-4 weeks in the planarian *Schmidtea mediterranea*
- Genotoxic stress does not prevent surviving stem cells from dividing but reduces their ability to increase proliferation rates and regenerate missing tissues after amputation.
- scRNAseq of wildtype vs. irradiated regeneration identified disrupted coordination of transient-regeneration activated cell states (TRACS), including elevated activin signaling
- Depletion of planarian *activin-2* rescues regenerative capacity, despite prior exposure to genotoxic stress and reduced stem cell numbers

## INTRODUCTION

Animals exhibit a remarkable diversity of regenerative capacities, from repair of individual organs to reconstruction of entire body plans. In many systems, adult stem cells support varying degrees of regeneration by proliferating and producing lineage-specific progenitors in response to injury^1,2^. The abundance, potency, and injury responsiveness of these populations are therefore important determinants of regenerative capacity^3–6^. However, stem cell activity is also regulated by differentiated tissues that provide instructive and supportive signals during tissue maintenance and regeneration^7–12^. Changes in these signals could therefore alter regenerative capacity independently of the number or potential of resident stem cells. Understanding how stem cell availability and tissue signaling together determine regenerative outcomes is essential to explaining both the robustness and limits of animal regeneration.

Because regeneration requires the ability of multiple cell populations to coordinate responses to tissue loss, sources of stress affecting any of these populations could disrupt the robustness and fidelity of repair. Infection^13^, endocrine stress^14^, and metabolic dysfunction^15^ have all been linked to reductions in regenerative capacity. Whether these conditions impair regenerative cells directly or alter the signals that regulate their activity is often difficult to resolve. Genotoxic stress presents a particularly revealing challenge because it can both deplete stem cells and affect the differentiated tissues surrounding them^16–18^. Failed regeneration could therefore reflect insufficient stem cell numbers, impaired responses within surviving stem cells, or disruption of the signaling network that normally coordinates tissue replacement. Determining where this coordination fails is essential to understanding how stress limits regenerative capacity and whether that inhibition can be reversed.

The freshwater planarian *Schmidtea mediterranea* provides a tractable system for examining how stress alters coordination between stem cells and differentiated tissues during whole-body regeneration. Planarian adult stem cells, called neoblasts, continuously divide and generate lineage-specific progenitors that maintain tissues throughout the animal^19^. Following injury, neoblasts exhibit an early proliferative response, followed by a second increase in proliferation associated with missing tissue^2,20,21^. This distinction allows ongoing stem cell division and population expansion to be examined separately from the proliferative response required for rapid tissue replacement. Amputation also induces transient regeneration-activated cell states (TRACS) in differentiated muscle, epidermal, and intestinal cells, including cells distant from the wound^16^. These states contribute to tissue patterning, stem cell proliferation, and remodeling, providing defined responses through which to examine coordination across the regenerating animal.

Despite this extraordinary regenerative capacity, planarians remain sensitive to physiological conditions that compromise tissue replacement. Pathogenic shifts in endogenous microbiota impair regeneration through activation of innate immune signaling, demonstrating that interactions between tissue condition and regenerative responses can limit repair^13^. Sub-lethal dosages of ionizing radiation are also a well-established tool for studying stem cell fate choices and differentiation, allowing the recovery of stem cell abundance to be followed over time^19,22,23^. Despite their potential to build and maintain tissues lineages, previous work showed that tissue fragments from sub-lethally irradiated animals could could not regenerate missing tissues^16^. However, whether this failure reflects insufficient numbers or impaired function of regenerative cells, altered wound-induced signaling in differentiated tissues, or both, remains unresolved. Comparing stem cell recovery with the activation of regenerative responses therefore provides a way to identify where coordination fails after genotoxic stress.

Here, we used low-dose genotoxic stress to distinguish the contributions of stem cell depletion and tissue signaling to regeneration failure in *S. mediterranea*. Our results identify Activin-dependent signaling as a reversible inhibitor of regeneration and reveal that altering the stress-induced signaling environment can restore tissue replacement despite stem cell depletion.

## RESULTS

### Relatively low doses of genotoxic stress inhibit planarian regeneration

Prior studies have shown that planarian regeneration is inhibited following a 1250R sub-lethal dose of ionizing radiation. To test the effect of underlying tissue and stem cell damage level on planarian regeneration, we first exposed planarian flatworms to increasing sub-lethal doses of ionizing radiation. Intact, healthy worms were irradiated with 250–1250 Rads and allowed to recover for one week before being amputated into head, trunk, and tail fragments (Fig. 1A). This interval was selected to allow time for the initial DNA damage response and stem cell depletion to occur. Unsurprisingly, all doses of ionizing radiation tested had subtle, but significant, effects on the number of proliferating stem cells in the animal, with 500R to 1250R doses resulting in a visible stem cell depletion (Fig. 1B, C). However, none of the tested doses resulted in a complete loss of stem cells or proliferation. Despite the presence of *piwi-1*-positive stem cells present prior to amputation, all tested dosages of irradiation produced regeneration defects, with near complete inhibition of regeneration with 750, 1000, or 1250 Rads (Fig. 1D, E). Phenotypes ranged in severity from cyclopia to no blastema formation. Lower doses of radiation yielded more animals with cyclopia and small blastemas, while higher doses – 750R and above –resulted in almost all fragments lacking blastema tissue at 14 days post amputation (Fig. 1E). Notably, inhibition of regeneration applied to both anterior and posterior facing blastema tissue, though regeneration of anterior structures was slightly more robust following irradiation in trunk fragments than in tail fragments (Fig. S1A-C).

**Figure 1.**
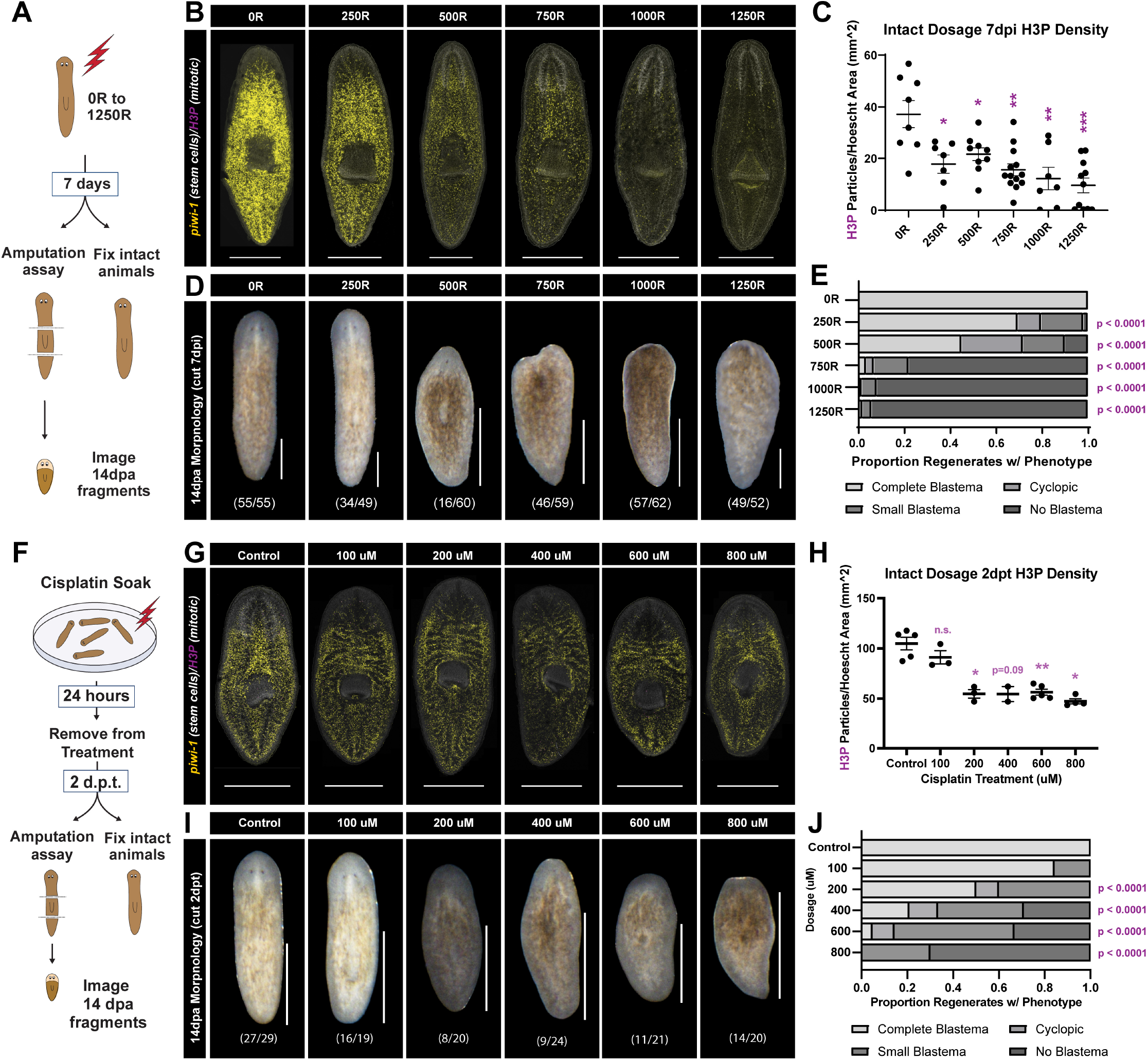
Low-dose genotoxic stress inhibits regeneration in *S. mediterranea.* (A) Schematic of irradiation dosage experiment (B) *piwi-1* in situ hybridization and phospho-histone H3 (H3P) antibody staining 7 days post 250R to 1250R (C) Quantitation of H3P density after 250R to 1250R with SEM bars 7 days post irradiation (dpi) (D) Morphologies of irradiated tail regenerates 14 days post amputation (14dpa) (E) Quantitation of irradiated tail regeneration phenotypes in regenerates 14dpa (F) Schematic of cisplatin dosage experiment (G) *piwi-1* in situ hybridization merged with H3P antibody staining 2 days post treatment (dpt) with cisplatin (H) Quantitation of H3P density with SEM bars 2dpt with cisplatin (I) Morphologies of cisplatin-treated tail regenerates 14dpa (J) Quantitation of cisplatin-treated regeneration phenotypes in regenerates 14dpa . P-values are Mann-Whitney test with correction for multiple comparisons. Scale = 1mm.

To test if inhibition of regenerative capacity was specific to low doses of ionizing radiation, or was a broader response to genotoxic stress, we soaked the animals in increases doses of the chemotherapy agent Cisplatin (Fig. 1F). All doses above 100μM resulted in reduced stem cell proliferation prior to amputation, with minimal observable depletion of stem cells (Fig. 1G, H). However, despite an abundant population of stem cells, cisplatin treated animals displayed significant and dose-dependent inhibition of regenerative capacity (Fig. 1I, J). Like regeneration after low doses of ionizing radiation, phenotypes ranged from cyclopia to full inhibition of blastema formation, with increasing doses showing increased severity of regeneration defects (Fig. 1J, Fig. S1D-F). Together, these experiments demonstrate that damage can significantly inhibit regeneration, even when proliferating stem cells remain.

### Inhibition of regenerative capacity is transient and can be rescued by re-injury

Given the rough correlation between regenerative capacity and the number of proliferating stem cells in the animal, we next wanted to determine if changing the recovery time after treatment would impact regeneration. To test this, worms were treated with either 750R or 1250R of ionizing radiation, then cut at 2, 7, 14, or 21 days after treatment (Fig. 2A, Fig. S2A). As expected based on prior research^23^, proliferating stem cell numbers in intact animals were initially depleted at 2, 7, and 14 days post irradiation but increased towards wild type numbers by 21 days post irradiation (Fig. 2B, C, Fig. S2B, C). Upon amputation, we found that regenerative capacity declined along with stem cell numbers, then began to recover at 14 to 21 days post irradiation, with peak inhibition of regenerative capacity at 7 days post irradiation (Fig. 2D, E). This trend was consistent across both anterior and posterior facing wounds in heads, trunks, and tails (Fig. S2D-J). Notably, the robustness of planarian regenerative capacity had not returned to the levels of unirradiated animals by 21 days post irradiation treatment, despite substantial numbers of resident adult stem cells. Together, these experiments indicate that sub-lethal doses of ionizing radiation can transiently inhibit regenerative capacity, despite a population of actively dividing and expanding stem cells.

**Figure 2.**
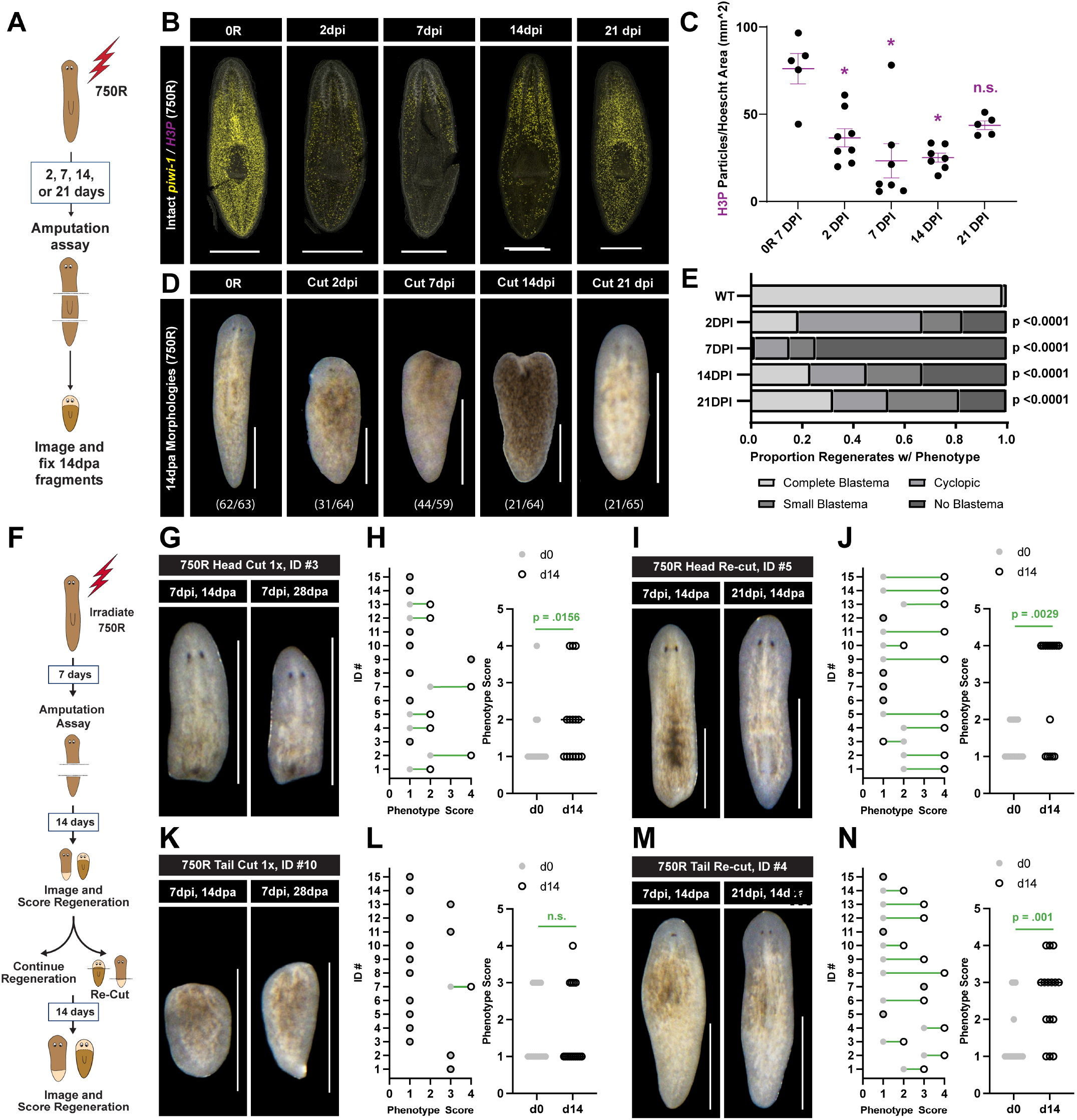
Inhibition of regeneration is transient and can be rescued by re-injury. (A) Schematic of irradiation time course experiment (750R) (B) *piwi-1* in situ hybridization merged with H3P antibody staining at 0, 2, 7, 14, and 21dpi (C) Scatter plot of H3P density quantitation with SEM bars at 0, 2, 7, 14, and 21dpi (D) Morphologies of tail regenerates at 14dpa (E) Quantitation of tail regeneration phenotypes in regenerates at 0, 2, 7, 14, and 21dpi (F) Schematic of re-cut experiment. (G) Head regeneration morphologies of head #3 cut 7dpi, imaged at 14dpa (left), and imaged again at 28dpa (right) (H) Scatter plot of individual head phenotype scores cut 7dpi, quantified at 14dpa (d0) and 28dpa (d14) (left), and summary of scores (right) (I) Head regeneration morphologies of head #5 cut 7dpi, imaged at 14dpa (left), then cut again at 21dpi, imaged 14dpa (right) (J) Scatter plot of individual head phenotype scores cut 7dpi, quantified at 14dpa (d0) and cut again at 21 dpi, quantified 14dpa (d14) (left), and summary of scores (right) (K) Tail regeneration morphologies of tail #10 cut 7dpi, imaged at 14dpa (left), and imaged again at 28dpa (right) (L) Scatter plot of individual tail phenotype scores cut 7dpi, quantified at 14dpa (d0) and 28dpa (d14) (left), and summary of scores (right) (M) Tail regeneration morphologies of tail #4 cut 7dpi, maged at 14dpa (left), then cut again at 21dpi, imaged 14dpa (right) (N) Scatter plot of individual tail phenotype scores cut 7dpi, quantified at 14dpa (d0) and cut again at 21 dpi, quantified 14dpa (d14) (left), and summary of scores (right). P values are Mann-Whitney test with correction for multiple comparisons. Scale = 1mm.

We next wanted to know of the recovery of regenerative capacity required an intact animal. In other words, if we were to re-injure a worm that had failed to regenerate after irradiation, would the secondary injury successfully launch regeneration. Or, does a failed regeneration event permanently inhibit regeneration. To address this question, we compared two injury paradigms. First, we cut heads, trunks, and tails at 7 days post treatment with either 750R or 1250R and measured regeneration phenotypes at 14 and 28 days post regeneration. In the second paradigm, regenerating fragments were re-injured with a transverse cut 14 days post amputation (21 days post irradiation) and then allowed to attempt to regenerate for 14 days. In both paradigms, regeneration phenotypes were scored at 14 days post amputation and again 14 days later with or without re-injury (Fig. 2F). In regenerates taken from animals 7 days post treatment with 750R, an additional 14 days of regeneration time slightly improved the phenotypes of head fragments regenerating tails, with some animals showing significant blastema growth (Fig. 2G, H). Re-injury at 14 days post amputation produced an even more dramatic phenotype improvement, with most heads regenerating a full posterior blastema and pharynx (Fig. 2I, J). In contrast, in tail fragments regenerating head tissue, additional time did not significantly improve phenotypes or blastema size (Fig. 2K, L). However, re-injury at 14 days post amputation resulted in significant improvement in phenotypes two weeks later, with most animals building anterior brain tissues (cyclopia or complete regeneration) (Fig. 2M, N). Similar results were achieved in fragments taken from animals treated with 1250R (Fig. S3I-P). Together, our results indicate that allowing the tissue to recover for a full three weeks after irradiation, in both whole worms and amputated tissue fragments, restores regenerative capacity in tissues treated with genotoxic stress. Notably, the re-injury experiment revealed that tail tissues, but not anterior brain structures, can eventually regenerate given enough time.

### Irradiation prevents the stem cell missing tissue response, but not proliferation or progenitor production

Successful regeneration in planaria depends on the coordinated activities of differentiated tissues that provide instructive and supportive cues, as well as proliferating stem cells that re-build missing tissues^2,24,25^. For example, amputation results in the induction of transient regeneration activated cell states (TRACS) in muscle, epidermal, and intestinal cell states, which are required for proper tissue polarity, stem cell proliferation, and tissue remodeling^26^. Downstream of TRACS, planarian stem cells engage in two proliferative bursts, one at 6 hours post amputation that is seen following all injuries, and one at 48 hours post amputation that is seen only when missing tissue regenerates^27^. Following these proliferative bursts, stem cell progenitors differentiate into missing tissues and support whole-body regeneration.

Since the regenerative capacity of animals after genotoxic stress roughly correlated with the number of proliferating stem cells present in the worm, we hypothesized that a rare stem cell sub-type might be required for regenerative capacity, resulting in failed regeneration when stem cells were not abundant. We decided to leverage our published scRNAseq atlas of planarian regeneration to compare stem cell states present in regenerating fragments taken from un-irradiated and sub-lethally irradiated (1250R) worms. To generate this dataset, worms were treated with either 0R, 1250R, or 6000R of ionizing radiation, amputated 3 days post irradiation, and regenerating biopsies were profiled with split-pool-ligation based RNA sequencing at 0, 1, 2, 4, 7, 10, and 14 days post amputation^26^. For our re-analysis, we first subsetted high-quality cells from only the 0R and 1250R time courses, re-normalized and re-clustered the dataset, and assigned tissues to broad lineage classes (Fig. 3A). We next identified all the stem cell and progenitor cells in both datasets using expression of known markers^22,28^ (Fig. 3B) and subset these cells into a separate object for re-normalization, clustering, and proportion analysis. Within the stem cell and progenitor subset, we readily identified progenitors biased towards epidermal, muscle, intestinal/phagocytic, secretory, and neural cell fates^28,29^ (Fig. 3C). We then split the 0R and 1250R time courses into their separate treatments (Fig. 3D) and analyzed the proportion of all captured cells that were stem cells and the proportion of stem cells distributed across the stem cell subtypes throughout regeneration (Fig. 3E). During regeneration in fragments taken from unirradiated animals, stem cells increased from 15-25% of captured cells. At 0 days post amputation, stem cells were distributed roughly equally across *piwi-1* high pluripotent stem cell states and most of the identified progenitor pools, with secretory progenitors being somewhat rare. These proportions were largely maintained at early timepoints (1-2dpa). At middle timepoints (4-7dpa), neural progenitors notably expanded at the expense of epidermal and intestinal/phagocytic progenitors, roughly correlating with the timeline for brain and head regeneration. At later timepoints (10-14dpa), neural progenitors remained enriched, with epidermal progenitor numbers recovering, and intestinal/phagocytic progenitors remaining rarer. In fragments taken from irradiated animals, however, stem cells were only 1% of captured cells at early timepoints, rising slowly to 5% by the middle of regeneration and 10% by the late timepoints. Prior to stem cell expansion, *piwi-1* high pluripotent stem cells and secretory progeny were not captured. However, by middle and late timepoints during regeneration, all stem cell sub-types had recovered and there were abundant *piwi-1* high pluripotent stem cells and neural progenitors present. Since we amputated irradiated animals a full 7 days after treatment with ionizing radiation (equivalent to 4dpa in the published dataset), the presence of all stem cell subtypes at that timepoint conflicted with our hypothesis that a particular stem cell state was missing in irradiated animals.

**Figure 3.**
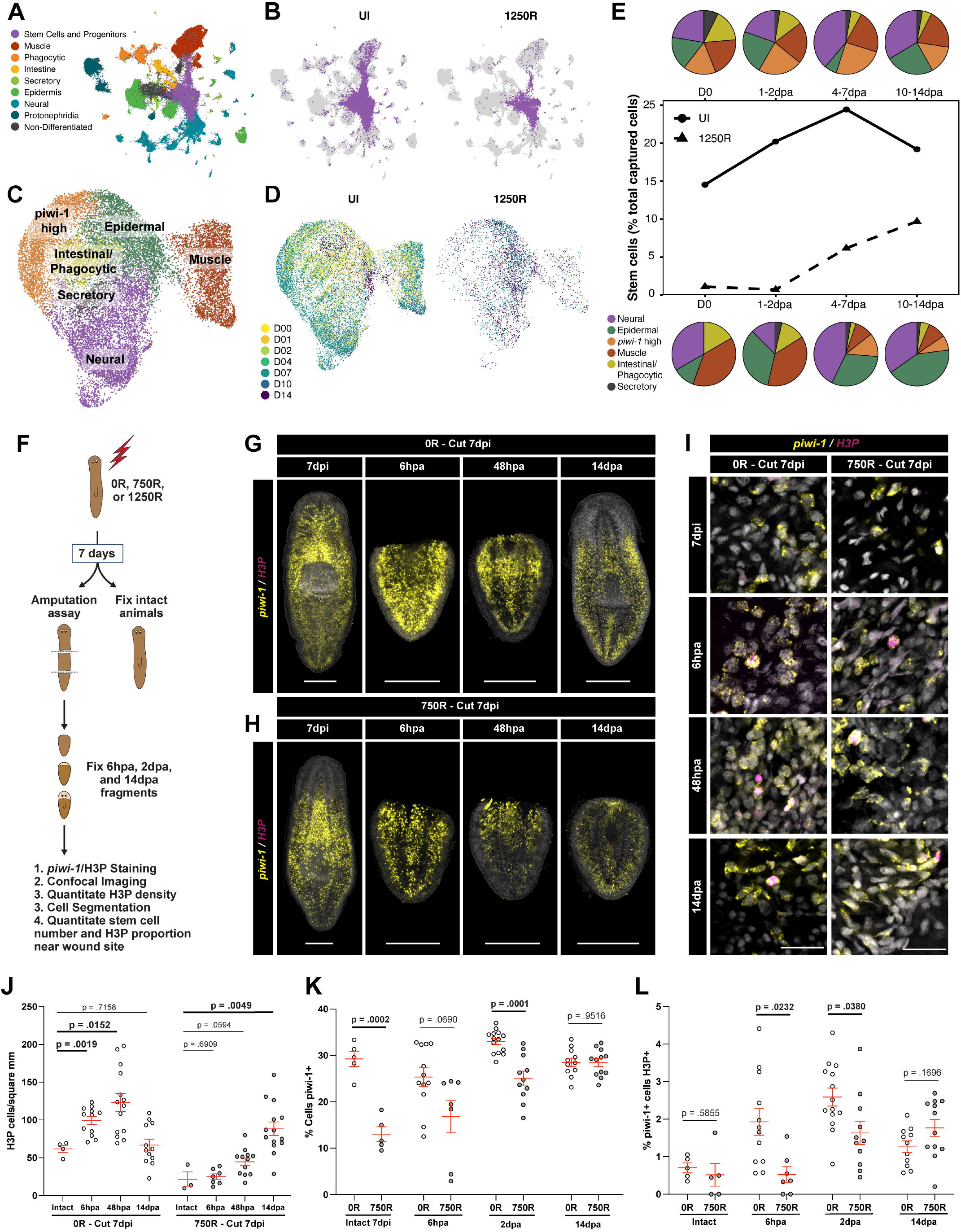
Reduced regenerative capacity is associated with loss of stem cell missing tissue response, rather than an inability to divide or produce progenitors. (A) Global UMAP embedding of unirradiated and 1250R regeneration time course dataset, labeled by tissue type. (B) Global UMAP embeddings highlighting stem cell and progenitor tissues in unirradiated and 1250R worms. (C) UMAP embedding of stem cell and progenitor subset, colored by stem cell sub-type (D) UMAP embeddings of stem cell and progenitor subset in unirradiated and 1250R worms, colored by regeneration timepoint. (E) Plot of stem cell proportion through time in unirradiated (solid line) and 1250R (dashed line) regeneration time courses. Circle plots depict the proportion of stem cells with stem cell sub-type identity at each phase of regeneration in unirradiated (above) and 1250R-treated (below) fragments. (F) Schematic of rradiation experiment followed by stem cell and H3P analysis. Whole mount max projections of intact and regenerating animals 6hpa, 48hpa, or 14dpa stained with *piwi-1* and H3P if cut 7dpi with 0R (G), or 750R (H). (I) Confocal slices of ROIs from animals G and H showing *piwi-1* and H3P staining. Quantitation of H3P density (J), percentage of cells *piwi-1^+^*(K) and percentage of *piwi-1^+^* cells also H3P^+^ (L) in animals from G and H. P values are Mann-Whitney test with correction for multiple comparisons. Scale = 1mm (G, H) or 25um (I).

We next turned to the hypothesis that the ability of the stem cells to respond to injury, the missing tissue response, was blocked by irradiation. To test this hypothesis, we quantitated stem cell numbers and proliferation rates after amputation in planaria treated with 0R, 750R, or 1250R of ionizing radiation. Following irradiation treatment animals were either fixed or amputated at seven days post irradiation (7dpi) and regenerating fragments were fixed at 6 hours, 2 days, or 14 days post amputation. Following fixation, tissues were stained with stem cells (*piwi-1*) and a dividing cell (phospho-H3) marker (Fig. 3F). Stem cells initially appeared partially depleted in irradiated samples relative to controls, but then returned to relatively normal densities by 14 days post amputation (Fig. 3G, H, Fig. S4A). Interestingly, stem cells accumulated near the wound site at 2 days post amputation in regenerating fragments taken from irradiated animals, despite subsequent failed regeneration. To measure the degree to which regenerating fragments could activate the early (6hpa) and late (2dpa) proliferative bursts, global H3P density was quantitated across all worms. While both proliferative bursts could be seen in unirradiated worms, mitotic activity exponential increased over time in the irradiated samples, reflective of stem cell expansion rather than regenerative stem cell behaviors (Fig. 3J, Fig. S4B). However, the reduction in mitotic cell counts could be due to depleted stem cell numbers, so we quantitated the percent of cells that were stem cells (*piwi-1^+^*) and the percent of stem cells in mitosis near the wound site. As predicted by observations of stem cells in whole animals, the percentage of *piwi-1^+^* cells was significantly depleted in both 750R and 1250R treated worms, but these numbers increased over time and returned to normal levels by 14 days post amputation (Fig. 3I, K, Fig. S4C, D). More importantly, the percent of stem cells dividing in intact worms seven days post irradiation was not significantly altered between unirradiated and irradiated worms, but the proliferative bursts at both 6 hours post amputation and 2 days post amputation were significantly reduced after irradiation (Fig. 3L, Fig S4E).

Together, these data strongly indicate that prior exposure to ionizing radiation does not prevent stem cell divisions, stem cell expansion, or generation of lineage-specific progenitors. Instead, irradiation inhibits the ability of stem cells to launch the ‘missing tissue response.’

### Irradiation disrupts coordination and patterning of transient regeneration activated cell states (TRACS) and elevated Activin signaling

The planarian missing tissue response is thought to be regulated primarily by wound-induced signaling in differentiated tissues. To investigate differentiated tissue responses to amputation after irradiation, we again leveraged our published single cell sequencing dataset, this time focusing on the differentiated tissues known to contain transient regeneration-activated cell states (TRACS). Previously, we identified TRACS in the muscle, epidermis, and intestine important for patterning, stem cell proliferation, and tissue remodeling during regeneration. To investigate whether TRACS still occurred following irradiation, we isolated and re-analyzed the muscle, epidermal, and intestinal cells from the unirradiated and post-1250R regeneration single cell time courses (Fig. 4A). In all cases, TRACS occurred in the irradiated samples, but with somewhat different frequencies. For example, *agat-1+* wound-induced epidermal cell states (E20) were noticeably reduced, while ira-1+ intestinal cells appeared somewhat increased (Fig. 4B) Visualization of TRACS *in vivo* confirmed that they all occurred in differentiated tissues following irradiation, but *agat-1*+ E20 cells had reduced frequency compared to unirradiated samples and *ira-1+* intestinal enterocytes were more plentiful (Fig. 4C, D). Analysis of TRACS cluster frequency as a proportion of the whole lineage over time in the un-irradiated and 1250R time course revealed more clearly how irradiation altered TRACS activation. *Notum*+ muscle cell states were muted, *agat-1*+ epidermal cell activation was prematurely truncated, and both intestinal cell TRACS were hyper-activated in irradiated worms, particularly the *ira-1*+ enterocyte cell state (Fig. 4E). In imaging of *notum+* muscle cells in irradiated worms, we also noticed a particularly striking phenotype, a loss of anterior pole formation (Fig. 4F, white arrow, Fig S5A), and the loss of polarized anterior notum expression (Fig. 4F, yellow arrow, Fig. S5B-D) in regenerates after ionizing radiation. Notably, both phenotypes have been previously observed in *follistatin* RNAi animals^30–32^. Returning to our scRNAseq dataset, we quantitated the percentage of cells expressing *follistatin* and it’s competing ligands *activin-1*, *activin-2*, and *activin-3* during regeneration in un-irradiated and 1250R-treated fragments. In addition, we quantitated the mean expression of *follistatin* and the Activin ligands within positive cells. This analysis revealed a consistent decrease in the number of *follistatin*+ cells and an increase in the number of cells expressing *activin-2* and *activin-3* (Fig. S5E), as well as increased activin expression levels of *activin-1* and *activin-2*, particularly during the later phases of regeneration (Fig. S5F). *In vivo* visualization of *follistatin* and *activin-2* confirmed a decrease in follistatin expression levels near the wound site (Fig. S5G, H), as well as increased expression of *activin-2* in the intestine (Fig. S5I). Together, these data indicate that low-dose irradiation prior to amputation generates an elevated activin signaling environment and disrupted TRACS coordination, ultimately resulting in a phenotype that closely phenocopies *follistatin* depletion.

**Figure 4.**
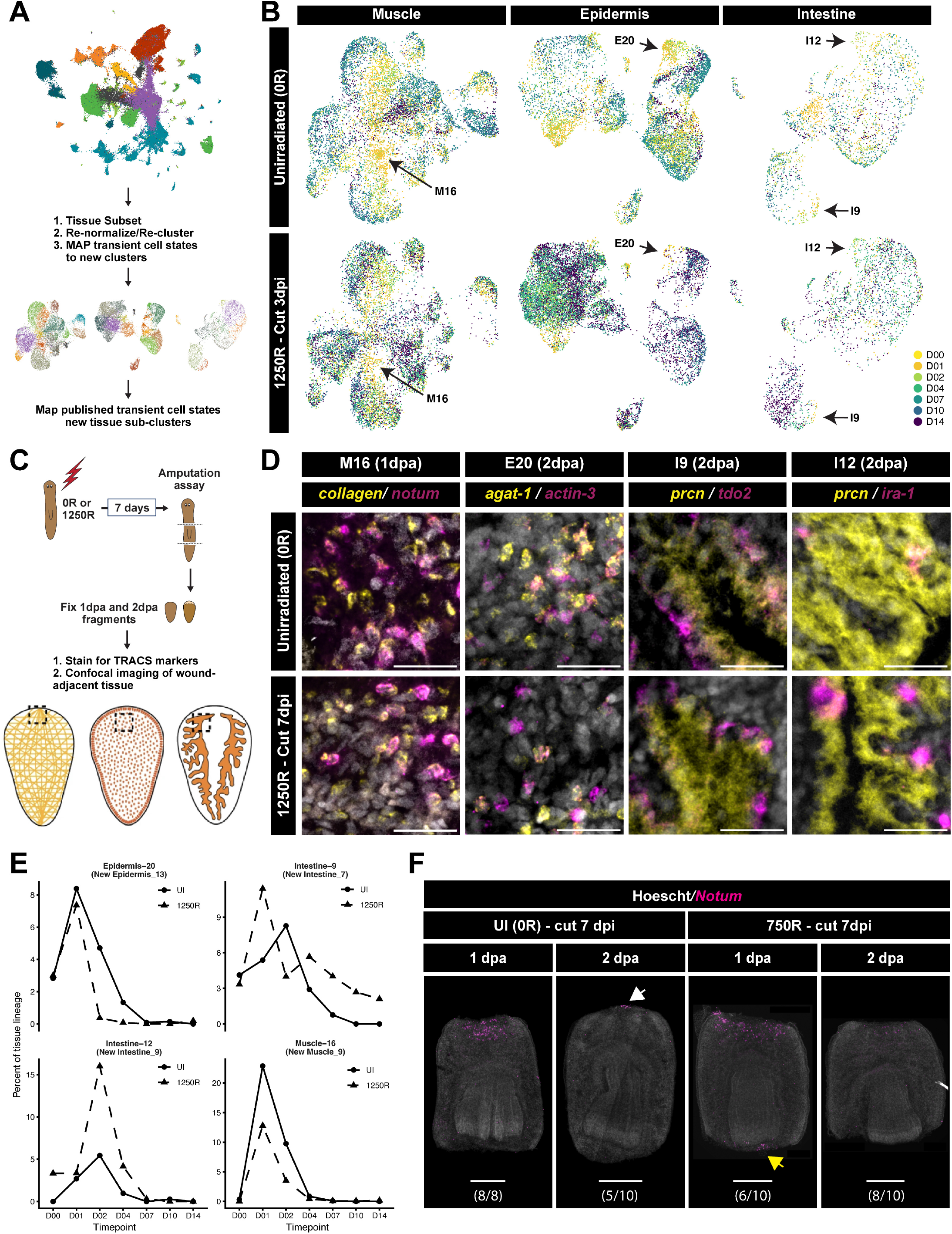
Regeneration after low-dose irradiation results in miscoordination of transient regeneration activated cell states. (A) Map published transient cell states to new tissue sub-clusters (B) UMAP embeddings of muscle, epidermal, and intestinal cells capture in Benham-Pyle et al. Nature Cell Biology 2021. Black arrows ndicate TRACS states that arise in both un-irradiated and irradiated biopsies during regeneration. (C) Depiction of TRACS staining protocol (D) Double FISH showing activation of TRACS genes (notum, actin-3, tdo2, ira-1) in muscle, epidermal, and intestinal cells (E) Proportion of cells in TRACS clusters through time n unirradiated (solid lines) and 1250R-treated (dashed lines) regeneration time courses. (F) Whole mount max projections of regenerating trunks cut at 7dpi fixed at 1 or 2dpa stained with notum.

### Inhibition of activin signaling rescues regenerative capacity after genotoxic stress

Activin/TGF-beta signaling is a highly conserved regulator of neural differentiation, stem cell proliferation, and activation of innate immune signaling pathways. Across multiple species, it has been linked to regulation of regeneration, tissue turnover, and cancer initiation, but with conflicting regulatory roles depending on the tissue or disease context^33–36^. Planaria possess three activin-like homologs, six activin-receptor-like homologs, and eight smad-like transcription factors. Given the observed increase in *activin-2* expression in regenerates exposed to low-dose ionizing radiation, we cloned all 18 Activin signaling homologs in *Schmidtea mediterranea* and screened them for a role in regeneration after irradiation using RNAi-mediated gene depletion. Worms were fed RNAi four times, irradiated with 750R, then cut into head, trunk, and tail fragments seven days post irradiation (7dpi). Regeneration phenotypes were scored 14 days post amputation (Fig. 5A). Of the 18 genes depleted, seven genes significantly altered regenerative capacity after irradiation in heads, trunks, or tails (Fig. 5B, Fig. S6). Notably, all three activin ligands improved regenerative capacity and the ability of tail fragments to regenerate anterior head structures by 14 days post amputation (Fig. 5B, C). In addition, one activin receptor (*ActR-4*) partially rescued anterior tissue regeneration, but with different penetrance in trunks and tails (Fig. 5B, C, Fig.S6). Of all the genes screened, only one produced worse regeneration outcomes, the transcription factor *smad4-1*, which resulted in tissue degeneration and, in some worms, lysis by 14 days post amputation in trunks (Fig. S6B, E). To find the genes with the strongest phenotypic rescues, we quantitated the size of the regenerated heads and pharynxes across all the RNAi conditions 14 days post amputation in regenerating tail fragments. *Activin-2* RNAi significantly increased the size of both the regenerated central nervous system and the regenerated pharynx (Fig. 5D, E). These results indicate that failure to regenerate after genotoxic stress may not be due to stem cell damage, but rather the active inhibition of regenerative capacity by elevated activin signaling.

**Figure 5.**
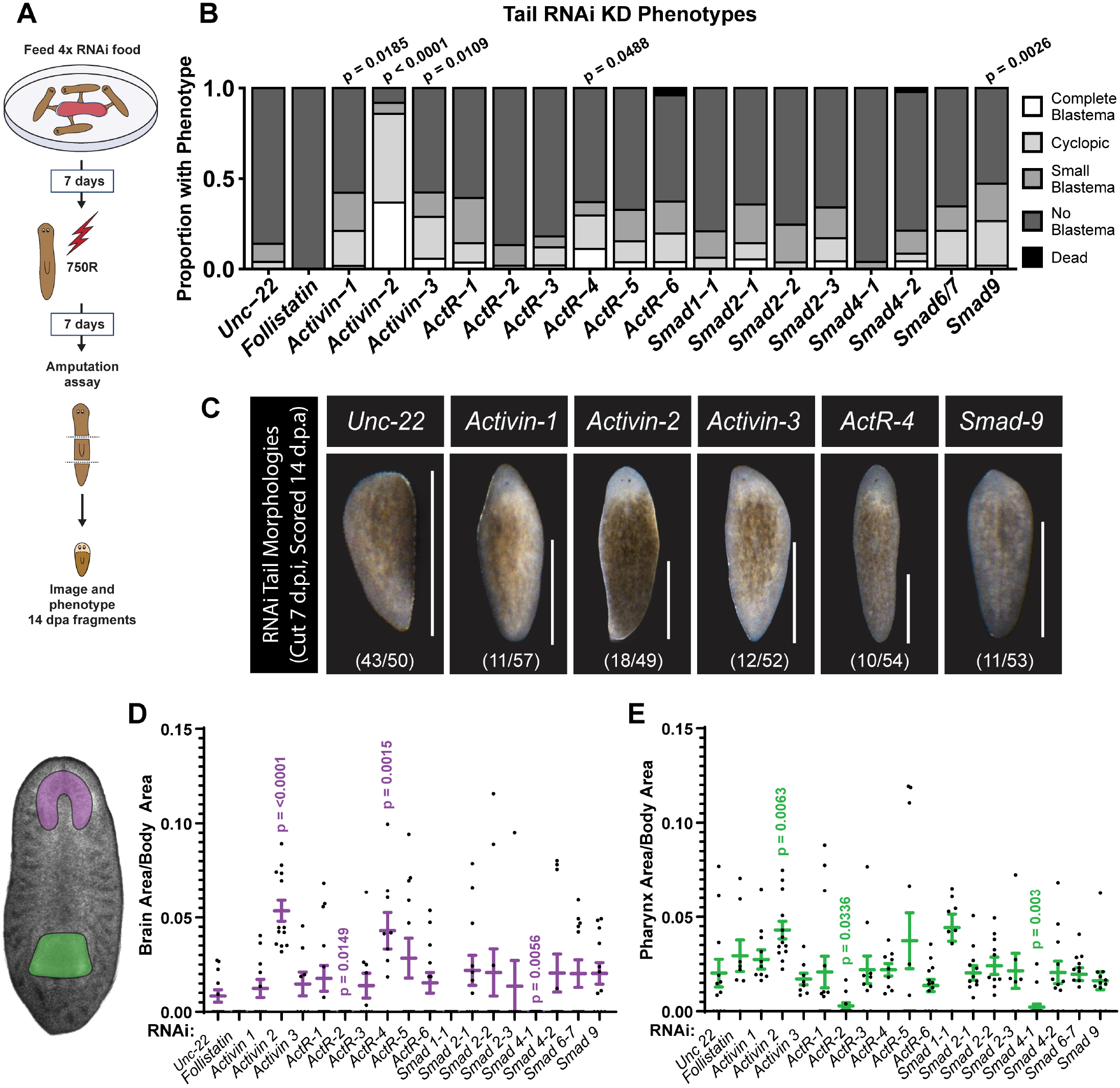
Activin signaling inhibits regenerative capacity after low dose irradiation. (A) Schematic of Activin homolog RNAi screen design (B) Phenotype quantitation of tail fragments 14dpa when cut from RNAi treated worms 7 days post 750R treatment (C) Morphologies of RNAi-treated tail regenerates with phenotypes distinct from Unc-22 controls. Quantitation of brain size (D) and pharynx size (E) in RNAi-treated tail regenerates 14dpa. P-values are Mann-Whitney test with correction for multiple comparisons. Scale = 1mm.

### Activin-2 depletion does not regulate stem cell loss after genotoxic stress, but rescues stem cell responses to amputation and brain regeneration

Prior studies have identified several signaling molecules in planarians that modulate stem cell resilience and regenerative capacity after irradiation. EGFR depletion results in a loss of proper symmetric divisions and an inability of stem cells to recovery after irradiation^23^, while depletion of ATM kinase was recently shown to increase stem cell survival and regenerative capacity following lethal radiation doses^37^. It has also been shown that beta-catenin depletion can rescue regeneration after mutagenic treatment^18^. To further dissect how planarian *activin-2* depletion restores regenerative capacity after irradiation, we quantitated stem cell numbers and proliferation rates after RNAi treatment, comparing intact animals 7 days after irradiation (7dpi), as well as regenerating fragments 6 hours, 2 days, and 14 days post amputation. We also compared regenerative stem cell responses in worms that had not been irradiated to determine if *activin-2* alters stem cell behaviors during normal regeneration (Fig. 6A).

**Figure 6.**
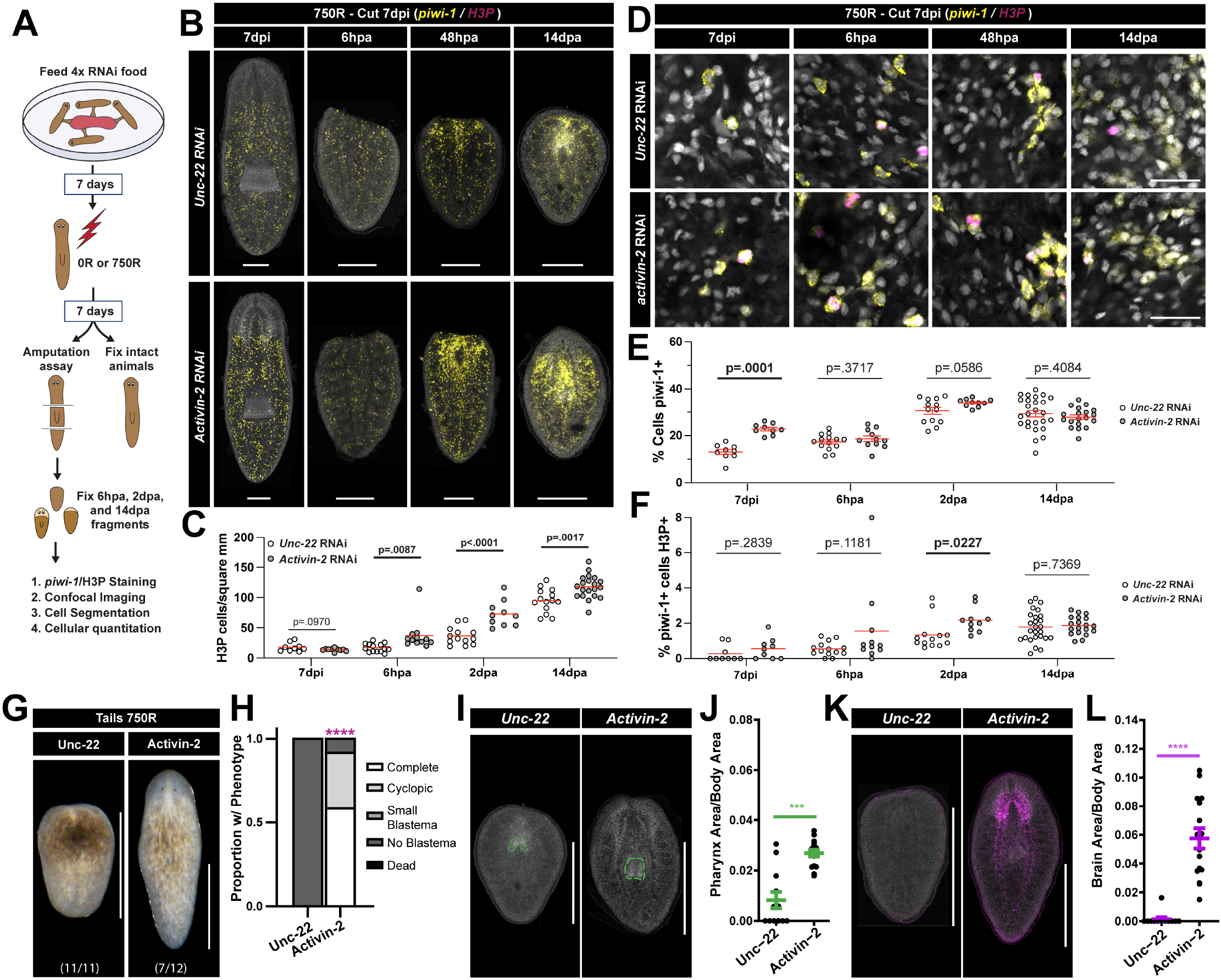
Activin-2 depletion rescues stem cell missing tissue response and regeneration of missing tissues. (A) Schematic of RNAi irradiation experiment followed by stem cell and H3P analysis. (B) Whole mount max projections of intact and regenerating animals 6hpa, 48hpa, or 14dpa, stained with *piwi-1* and H3P (C) Quantitation of H3P density in animals depicted in B. (D) Confocal slices of ROIs from animals in B, showing *piwi-1* and H3P staining. Quantitation of percentage of cells *piwi-1^+^* (E) and percentage of *piwi-1^+^* cells also H3P^+^ (F) in animals from B. P values are Mann-Whitney test with correction for multiple comparisons. (G) Morphologies of RNAi-treated tail regenerates (H) Phenotype quantitation of tail fragments 14dpa when cut from RNAi-treated worms 7 days post 750R treatment. (I) Representative mages of pharynx size in RNAi treated worms 14dpa when cut 7 days post 750R (J) Quantitation of normalized pharynx area for worms in I. (K) Representative images of brain area stained by PC-2 in situ hybridization in RNAi treated worms 14dpa when cut 7 days post 750R (L) Quantitation of brain size for worms in K. Scale = 1mm (B, G, I, K) or 25um (D)

Global proliferation levels in both unirradiated and irradiated worms were quantitated first. We observed no change in the normal induction of proliferation following amputation in un-irradiated animals treated with either control (*unc-22*) or *activin-2* RNAi, other than a slight, but significant decrease in mitotic density in intact worms (Fig. S7A, B). However, irradiated worms did show a significant difference in proliferation. While control worms displayed a slow increase in proliferation after amputation indicative of stem cell expansion, *activin-2* RNAi worms showed an increase in proliferation as early as 6 hours post amputation, with significantly higher proliferation than controls at 6 hours, 48 hours, and 14 days post amputation (Fig. 6B, C).Visualization of *piwi-1* positive stem cells also revealed more accumulation of stem cells near the wound site at 48 hours post amputation in *activin-2* RNAi animals compared to controls (Fig. 6B). We further quantitated the percentage of cells that were *piwi-1* positive and the percentage of stem cells dividing to understand how activin signaling might be regulating stem cell behaviors. In unirradiated worms, we found very few differences between control and *activin-2* RNAi worms. There was a small but significant increase in stem cell number in intact worms, as well as in worms two days post irradiation, and a slight decrease in proliferation rates at two days post amputation (Fig. S7C-E), but in both cases these changes were subtle and did not cause any change in regeneration outcome or tissue patterning. In irradiated worms, *activin-2* RNAi led to a significant increase in total stem cell numbers in intact worms at 7 days post irradiation, when amputations were performed (Fig. 6D, E). To determine why this might be the case, we analyzed stem cell numbers and global proliferation rates in control and *activin-2* RNAi treated worms after irradiation in the absence of an amputation (Fig. S7, F-O). Activin-2 depletion resulted in a slower loss of stem cells, but comparable numbers at 3 days post-irradiation (Fig. S7F, G, H). However, proliferation rates were higher at 2 and 7 days post irradiation (Fig. S7K, M, N), resulting in a higher number of stem cells by 10 days post amputation (Fig. S7F, J).

Interestingly, the difference in stem cell number prior to amputation that was present in control versus *activin-2* RNAi animals was not associated with increased stem cell proliferation rates. Moreover, there was no apparent difference in stem cell number in control vs. activin-2 RNAi animals at any time points after amputation (Fig. 6D-F). Indeed, the primary difference between control and *activin-2* RNAi worms’ responses during regeneration was that *activin-2* RNAi animals had significantly higher amputation-induced proliferation at 48 hours post amputation (Fig. 6F), the proliferative burst most associated with the ‘missing tissue response’ that drives rapid regeneration. The significance of this rescued missing tissue response was particularly evident when we compared the blastema growth, pharynx regeneration, and brain regeneration of head, trunk, and tail regenerates in control and *activin-2* RNAi treated animals. In heads, where pharyngeal progenitors are often present in the pre-existing tissue, irradiated fragments were unable to build a blastema but could often re-build a pharynx (Fig. S8A-F). Upon *activin-*2 depletion, blastema formation was rescued. In trunks, irradiated fragments were unable to build brains or posterior blastema tissue, both of which were rescued by *activin-2* RNAi (Fig. S8G-L). In tail fragments, irradiated regenerates were unable to build brain or anterior blastemas, and displayed delayed pharyngeal regeneration, all of which were rescued upon *activin-2* depletion (Fig. 6G-L).

To test whether increased regenerative capacity after *activin-2* depletion was specific to ionizing radiation stress, we treated RNAi-fed animals with 800uM Cisplatin, then amputated them into head, trunk, and tail fragments (Fig. S9). In all body regions, *activin-2* RNAi significantly improved regeneration phenotypes following Cisplatin exposure. In heads regenerating tail fragments, the number of animals able to build a posterior blastema increased from 0% to 50% (Fig. S9A, B). Trunks and tails demonstrated increased ability to build brain tissue and photoreceptors (Fig. S9E-F, I-J). As seen with ionizing radiation, *activin-2* depletion had no significant effect on regeneration in control animals (Fig. S9C-D, G-H, K, L).

Together, these data indicate that the rescue of regeneration in activin-depleted worms treated with ionizing radiation is due to an increased missing tissue response in stem cells and accelerated expansion post-amputation, rather than a dramatic change in the number of stem cells that survive the initial radiation dose. Moreover, this increased expansion and proliferation of stem cells rescues rapid blastema growth and regeneration of brain tissue. These results reveal that after ionizing radiation, stem cells are not incapable of regenerating missing tissue due to high levels of DNA damage or apoptosis. Instead, stem cells are being actively inhibited from regenerating missing tissue by an external cue – elevated Activin signaling.

## DISCUSSION

### Stem cell abundance and injury responsiveness make distinct contributions to regeneration

Following sublethal irradiation, stem cell abundance broadly correlated with regenerative capacity. Both declined after treatment and recovered over time. However, our findings indicate that this relationship does not reflect a simple dependence of regeneration on stem cell number. Surviving stem cells continued to divide, expand, and generate diverse lineage-biased progenitor states, yet they failed to mount normal amputation-induced proliferative responses and their progeny failed to differentiate into missing tissue. Moreover, *activin-2* depletion restored substantial tissue regeneration, despite persistent stem cell loss and comparable prior exposure to DNA damage. Thus, stem cell abundance and damage alone cannot explain regenerative outcomes after genotoxic stress. The signals regulating surviving stem cell function and responsiveness to amputation are also critical.

These observations may inform models of how planarian stem cell heterogeneity contributes to regenerative capacity. Although some stem cell states were not captured immediately after irradiation, all identified stem cell and progenitor classes were present at later time points when regeneration remained impaired. Persistent loss of a single indispensable subtype^20,38^ is therefore unlikely to explain the regeneration block. Instead, our findings are consistent with models in which regenerative potential is broadly distributed across planarian stem cell states and diverse progenitors can be produced through flexible or stochastic fate decisions^22,39,40^. While our experiments do not directly resolve the mechanisms of fate specification, they argue that the presence of diverse stem cell states does not, by itself, ensure a productive regenerative response.

Our findings, instead, indicate that whether surviving stem cells can regenerate depends on whether they respond to tissue loss. The rescue by *activin-2* depletion suggests that stem cell regenerative potential remains accessible, but is retrained by the signaling environment after genotoxic stress. One possibility is that elevated activin signaling interferes with their ability to respond to otherwise productive cues from transient regeneration-activated cell states (TRACS)^16,32^. Alternatively, stress may disrupt the coordination of those cues themselves. The success of re-injury after recovery is compatible with either possibility – a new injury may provide regenerative instructions once inhibitory signaling has diminished. Understanding how surviving stem cells respond therefore requires considering both the coordination of tissue derived signals and the environment in which those signals are received by stem cells.

### Regeneration depends on coordinated responses across tissues

Regeneration requires differentiated tissues to do more than recognize an injury. They must coordinate signals that direct stem cell activity, establish tissue polarity, and remodel existing structures as new tissues form^16,41,42^. Our findings suggest that genotoxic stress disrupts this coordination without eliminating underlying responses. Transient regeneration activated cell states (TRACS)^16^ remained detectable after irradiation, but their activation diverged across tissues: *notum+* muscle responses were muted, *agat+* cell activation was prematurely truncated, and intestinal responses were exaggerated. Failed regeneration was therefore accompanied by a mixture of diminished and excessive responses, suggesting that the balance among regeneration-associated states may be as important as their individual activation.

The altered *Activin/Follistatin* balance we observed in irradiated animals offers a potential connection between altered tissue responses and the failure of stem cells to regenerate. Increased activin expression and reduced wound-associated *follistatin* could create an environment in which regenerative instructions are produced but cannot elicit an effective response from stem cells^30,41^. Alternatively, elevated activin signaling may help drive the altered TRACS dynamics, disrupting the instructions themselves. Importantly, these possibilities are not mutually exclusive. Signals that restrain stem cell responsiveness to amputation could also alter the differentiated tissues responses needed to sustain regeneration. The rescue of regeneration by *activin-2* depletion identifies a critical cue in the signaling environment, but whether depletion restores TRACS coordination, releases stem cells from inhibition, or accomplishes both remains an important question.

Together, our observations indicate that miscoordination among differentiated tissues and stem cells could provide a framework for understanding regenerative failure more broadly. An animal may retain stem cells, produce progenitors, and activate regeneration-associated tissue states in surrounding differentiated tissues, yet still fail to regenerate missing tissues because these responses no longer work together. From this perspective, identifying which responses persist – and how their signaling relationships change – may be as revealing as identifying responses that are lost. Defining where coordination breaks down could expose opportunities to restore regeneration by bringing persistent cellular capacities back into cooperation.

### Changing the signaling environment can unlock regenerative capacity

The rescue of regeneration by *activin-2* depletion offers an encouraging implication – substantial regenerative capacity can remain accessible after damage. This possibility has precedents beyond planarians. In the mammalian brain, irradiation impairs neurogenesis through changes in both neural precursors and their environment^17^. Indeed, un-irradiated neural precursors transplanted into the irradiated hippocampus fail to differentiate into neurons, and inflammatory blockade augments neurogenesis after irradiation^43^. More directly paralleling our findings, vascular-derived TGF-beta increases in the irradiated mouse neural stem cell niche, and blocking this signaling improves neurogenesis^44^. Together, these studies and our results demonstrate that the consequences of genotoxic stress extend beyond the stem cells initially damaged to include tissue-derived signaling that continue to restrict new cell production.

The prospect of changing regenerative outcomes through focused signaling interventions has gained further support from comparative studies of regenerative and non-regenerative mammals. Macrophages are required for ear pinna regeneration in spiny mice, and factors secreted by activated spiny mouse macrophages promote regenerative behaviors in fibroblasts from both spiny mouse and *mus musculus* ^45,46^. Moreover, restoring deficient retinoic acid signaling reactivated ear pinna regeneration in mice, demonstrating that a restricted regenerative outcome can conceal a capacity for substantially greater tissue repair^47^. Our findings extend this principle to regeneration after genotoxic stress – depletion of a single ligand restored substantial regeneration despite prior irradiation or cisplatin exposure, with irradiation rescue occurring while stem cell numbers remained reduced. Although restoring an evolutionary limited response and overcoming damage-induced inhibition present different challenges, both reveal that changing a specific signaling input can profoundly alter what resident cells can accomplish.

An emerging model is therefore that augmenting regeneration requires identifying which signals are missing, which are excessive, and how these signals shape the behavior of cells that remain available for the repair. In some settings, supplying a regenerative instruction may be sufficient. In others, an inhibitory influence must be reduced before existing instructions can produce a response. *Activin-2* depletion provides evidence for this second opportunity. The encouraging prospect is that restoring regeneration after damage may not always require rebuilding the stem cell population first. Changing conditions under which surviving cells operate can allow them to achieve far more.

### Understanding and releasing restraints on regeneration

Moving forward, the strength of the *activin-2* rescue reported here raises an important mechanistic question: how does this ligand exert such a substantial influence over regenerative capacity? Although depletion of all three activin ligands individually improved regeneration outcomes, *activin-2* depletion produced the strongest rescue of regenerated brain and pharynx size. Differences in expression, protein structure, or receptor specificity could contribute to this effect. The weaker rescues following individual receptor depletion suggests additional complexity. *Activin-2* is expressed primarily in muscle and intestinal tissues, whereas its potential receptors are distributed across many cell types, including stem cells, parenchymal phagocytic cells, and neurons^16,31,48^. Moreover, activin signaling has been implicated in broader innate immune responses in planarians^49^. Ligand depletion may therefore alter several cooperating tissue responses, while receptor perturbations are buffered by redundancy or disrupt both supportive and inhibitory signals. Identifying the relevant receiving cells will help distinguish if activin-2 limits regeneration by altering tissue-derived instructions, restraining stem cell responses to those instructions, or both.

Beyond how activin-mediated restraint of regeneration operates lies the question of why regenerative restraint would arise at all in an organism. Why would a mechanism limiting regenerative ability be evolved or maintained? One possibility is that transient inhibition limits regeneration while damaged cells and tissue recover, thereby limiting inheritance of mutations or propagation of damage to regenerated tissues. Alternatively, elevated activin signaling may be a consequence of disrupted tissue physiology that interferes with regeneration without providing a protective benefit. Our experiments do not establish the purpose of regenerative inhibition, but its reversibility makes the distinction interesting. The recovery of regeneration after re-injury and its rescue by *activin-2* depletion show that an unsuccessful regenerative response need not reflect a permanent loss of regenerative ability. Determining what induces the inhibitory environment, what allows it to subside, and the functional outcome when inhibition is pre-maturely released will be essential to understanding whether it represents an adaptive response, a failure of coordination, or some combination of the two.

Our results and the questions they generate invite a broader view of what sets regenerative limits. The regenerative capacity an animal displays as it ages or after injury may fall well below what its surviving stem cells can accomplish under different signaling conditions. In our experiments, changing a single signaling input allowed a depleted stem cell population to meet a substantial regeneration challenge despite prior genotoxic stress. Moving forward, an important challenge will be to learn when and where such restrained potential exists and how to engage it^47,50,51^. Understanding the signals that hold regeneration in check may reveal that the limits imposed by damage or evolutionary history are far less fixed than they appear.

## RESOURCE AVAILABILITY

Further information and requests for resources and reagents should be directed to, and will be fulfilled by, the lead contact, Dr. Blair Benham-Pyle. This study generated RNAi feeding vectors containing cDNA inserts of Activin signaling components, as well as microscopy data and data analysis scripts. These vectors, data, and scripts will all be shared by the lead contact upon request following publication.

## ACKNOWLEDGEMENTS

We thank members of the CAGT Stem Cell and Regenerative Medicine Center and the Benham-Pyle Lab for valuable input into this project. This project was supported by the Cancer Prevention Research Institute of Texas (RR210037). H.B. was also supported by the BCM post-baccalaureate research education program (R25GM069234). Z.C. and A.R. were also supported by the Genetics and Genomics Training Program (T32GM139534). B.W.B-P. was also supported by Baylor College of Medicine and NIH DP2AG093210.

## AUTHOR CONTRIBUTIONS

H.B. performed experiments, analyzed data, made the figures, and wrote the paper. A.D. performed experiments, analyzed data, and made the figures. Z.C., N.M., A.R., and H.A. performed experiments and analyzed data, B.W.B-P designed the project, obtained funding, performed experiments, analyzed data, made the figures, and wrote the paper.

## DECLARATION OF INTERESTS

The authors declare no competing interests.

## MATERIALS AND METHODS

### Animal Husbandry

*S. mediterranea* animals from the asexual clonal strain CIW-4 (C4) were maintained in 1x Montjuic salts in a planarian recirculation culture system where they were fed chunk beef liver biweekly^13,52^. Prior to irradiation and RNAi treatments, planarians were pulled from the aquatics system and starved in static culture tupperware boxes at 20°C for at least 7 days before use. Following irradiation, cisplatin, and RNAi treatments, planarians were maintained in petri dishes in 1x Montjuic + gentamicin (50ug/ml).

### Amputations and Regeneration Phenotype Scoring

When performing amputations for regeneration studies, planarians were amputated into head, trunk, and tail fragments by cutting above and below the pharynx. At various days post-amputation (d.p.a), regeneration phenotypes were assigned a phenotype based on blastema size and patterning, which included scores of (0) dead/lesions, (1) no blastema formation, (2) small/incomplete blastema formation, (3) cyclopia, and (4) complete blastema formation. On the same day, live images were taken of regeneration phenotypes on a Nikon SMZ18 brightfield stereomicroscope. Trunk regeneration was scored based on the anterior-facing blastema.

### Cloning of S. mediterranea transcripts

The Sánchez Alvarado Lab reference transcriptome (smed_20140614) was used as the reference sequence for designing PCR primers to amplify transcripts of interest or to design cloning inserts for synthesis. Several Activin signaling molecules (Activin-2, Follistatin, ActR-1, ActR-3) had been previously cloning into RNAi feeding vectors by Gibson cloning as previously described^26^. For these, Primer3-optimized PCR primers were used to amplify target regions (500-1000bp) from the cDNA template and overhangs homologous to the pPR-T4P vector were added to the 5’ ends of each primer. PCR products were inserted into linearized vector using Gibson Assembly, then transformed directly into E. coli strain HT115. Insert sequences were verified by full plasmid sequencing. For newly cloned genes, inserts were selected from reference cDNA and flanked by T7 and primer/polymerase binding sites optimized for dsRNA and probe synthesis^53^. The combined sequence was synthesized and inserted into by Twist Biosciences. SMEDIDs for all cloned constructs are listed in Table 1.

### RNAi Food Preparation and Feeding

RNAi feeding vectors were transformed into the HT115(DE3) *E. coli* strain and stored in glycerol stocks at −70℃ until RNAi food preparation. For RNAi food preparation, the bacteria containing our plasmids of interest were inoculated in 2XYT media containing Kanamycin (50ug/mL) and Tetracycline (50ug/mL) and incubated at 37℃ with shaking for 16 hours. For high throughput preparations, bacterial glycerol stocks were inoculated into 2mL of media in a 24-well, 10mL deep well plate. For normal preparations, bacteria were inoculated into 8mL of media in 50mL BioReactor conical tubes. After incubation, additional 2XYT media with Kanamycin (50ug/mL), Tetracycline (50ug/mL), and IPTG (1mM) was added to the cultures, 6mL for high throughput and 24mL for normal preparations. The cultures were then placed back into the incubator for 4 hours to allow for induction of dsRNA by IPTG. After induction, bacteria were spun down for 10 minutes at 3600 rpm. Beef liver puree was mixed with food coloring, then the RNAi bacterial pellet was resuspended in the liver and stored at −20℃. Planarians were fed RNAi liver four times over the course of two weeks, with one to two days in between each feeding. RNAi fed planarians were kept in petri dishes in 1x Montjuic + gentamicin (50ug/ml), replacing the water after each feeding, and fresh dishes after every 2 feedings.

### *In situ* hybridization and antibody staining

DIG- and FLCN-conjugated probes were synthesized as previously described^54^. Samples were fixed using either NAFA^55^ or NAC^54^ fixation protocols. NAFA fixation was used for single FISH and antibody stained worms, while NAC fixation was used for *PC-2* and double FISH stained worms. Following fixation, samples were stored in methanol at −20C until use. In situ hybridization was performed as previously described^16,54,56^. In brief, samples were bleached for 3 hours using a modified Ryan King’s formamide bleach, washed, and incubated in a proteinase K/SDS solution, followed by post-fixation (4% PFA, 0.3% Tween), then hybridized overnight with a RNA probes at 56C (DIG and/or FLCN conjugated). The following day, samples were washed through a series of wash buffers at 56C (Wash Hybe, 2X SSC, .2X SSC) then transition into MABT block (5% horse serum, 0.5% Roche Western blocking reagent in MABT), then incubated overnight at 4C with anti-DIG-POD antibody (1:1000; SIGMA 11207733910). Probes were developed using either FLCN- or Cy3-tyramide (1:2000 or 1:1000, respectively) in borate buffer with 0.0006% H2O2 (1:5000) for 45 minutes. For double FISH, following DIG-probe development, samples were incubated in peroxidase (200mM NaN3 in PBSTw) for 1-2 hours, then incubated overnight at 4C with anti-FLCN-POD antibody (1:1000; SIGMA 11426346910) and Hoechst (1:500). The FLCN probe was developed using FLCN-tyramide (1:1000). For antibody staining, samples were blocked following probe development (5% horse serum, 0.5% Roche Western blocking reagent, 0.5% goat serum in PBSTx) for 2 hours, then incubated overnight at 4C with primary antibody (rabbit anti-H3P, 1:500; abcam Ab32107). The next day, samples were washed with PBSTx, then incubated overnight in secondary antibody + Hoescht (anti-rabbit Alexa 555, 1:1000; abcam ab150086). Upon completion of staining protocols, samples were post-fixed in 4% PFA for 1 hour, then cleared at least overnight in DAPCO/ScaleA2.

### Microscopy

Confocal images were captured on a Zeiss confocal microscope using the Zen software package. Live brightfield images were captured on a Nikon SMZ18 Manual Digital Sight 10, using NIS-Elements Discover software. Epifluorescent images were taken on a Nikon SMZ25 DS Qi2, using the NIS-Elements Advanced Research software.

### Image Analysis

H3P density was calculated as previously described^16,56^. In brief, a custom Fiji Script was used to max project, threshold, and binarize the hoescht and H3P channels of whole worm confocal tiled z-stacks to generate whole worm and H3P+ cellular masks. H3P+ cells were then counted and normalized by whole worm area to generate an H3P density for each worm. To quantitate the percentage of *piwi-1*+ and H3P+ cells in each worm, we used the multichannel-nuclear- analysis fiji macro freely available from Ariel Waisman (https://multichannel-nuclear-analysis.github.io/) with minor modificiations. In brief, 20X ROI zstacks were collected from the post-pharyngeal region of intact worms or within 100 microns of the wound site in amputated animals. A custom Fiji script was used to normalize the intensity distributions of the zstack to account for reduced light penetration, then split each zstack into individual images. The collection of images for each worm was processed by the multichannel nuclear analysis macro to segment nuclei and calculate the mean intensity of *piwi-1* and H3P per cell. Thresholds for piwi-1 and H3P positivity were set as a multiplier of the channel background detected by the nuclear analysis macro (1.5X background for *piwi-1*, 4.5X background for H3P). Summary statistics pooling all cellular data from each worm were generated using a custom R script, with the final output producing the total number of cells analyzed per worm ROI, the total number of piwi-1+ cells, the total number of H3P positive cells, and the % of cells piwi-1+, H3P+, or double positive. All custom scripts are readily available by request from the corresponding author upon publication.

### Analysis of published scRNAseq datasets

To examine cell-state composition and TRACS prevalence following sublethal irradiation, we reanalyzed published scRNA-seq data from the unirradiated and 1250R regeneration timecourse in Benham-Pyle et al. Nature Cell Biology 2021^16^. Cells from these treatments were selected from the published Seurat object, and a new object was constructed from raw RNA counts while retaining sample, treatment, time-point, and published tissue and cluster annotations. Analyses included days 0, 1, 2, 4, 7, 10, and 14 post amputation. Cells with fewer than 1,000 RNA counts or 200 detected genes were excluded. Retained cells were normalized using SCTransform, followed by principal component analysis (PCA). Global UMAP embeddings and nearest-neighbor graphs were generated using the first 150 principal components. Two- and three-dimensional UMAPs used a minimum distance of 0.2, and global clusters were identified using Seurat’s FindClusters function with default parameters. The recorded analysis environment comprised R 4.6.1 and Seurat 5.5.1.

Tissue identities were assigned using reference-based label transfer and cluster-enriched genes. An annotated reference object containing 2,000 cells per sample from the published dataset was normalized using SCTransform. Transfer anchors were identified using SCT normalization with 50 principal components, and tissue labels and prediction scores were transferred using dimensions 1–50. Predicted tissue labels were generated with a prediction-score filter of 0.15. Global clusters were manually assigned to nine tissue groups using tissue predictions and marker expression: stem cells and progenitors, secretory, protonephridia, phagocytic, neural, muscle, intestine, epidermis, and non-differentiated cells. Global clusters 4 and 19 were assigned to the non-differentiated group.

Each tissue group was reconstructed from raw RNA counts and independently normalized using SCTransform, followed by PCA, UMAP, nearest-neighbor graph construction, and clustering. Tissue-specific analyses used 15 principal components for phagocytic, intestinal, protonephridial, and stem/progenitor cells; 20 for epidermal, muscle, secretory, and non-differentiated cells; and 45 for neural cells. Clustering resolution was 0.6 except for stem/progenitor cells (0.4) and neural cells (0.7). Stem/progenitor clusters 1, 4, and 7 were subsequently combined, yielding six final subclusters. Updated tissue-specific subcluster assignments were transferred to the global object. Correspondence between published and updated clusters was quantified by shared cell membership, expressed as fractions of both the original and updated clusters. Global and tissue-specific UMAPs were visualized by tissue, subcluster, treatment, and time point.

Cell coverage was tabulated for each tissue and subcluster by treatment and time point, with absent subclusters recorded as zero cells. Coverage categories were fewer than 50, 50–99, and at least 100 cells. Temporal coverage eligibility was recorded for populations containing at least 300 cells per treatment and at least 20 cells at five or more time points; matched eligibility required these criteria in both treatments. These coverage flags were reported separately from the occupancy calculations.

Stem/progenitor abundance was calculated as the percentage of all captured cells at each treatment and time point. Time points were grouped into baseline (day 0), early (days 1–2), middle (days 4–7), and late (days 10–14) phases. Phase-level abundance was summarized as the arithmetic mean of the constituent time-point percentages. Stem/progenitor subtype composition was calculated from pooled cell counts within each treatment and phase.

Focused TRACS analyses examined the published Epidermis-2, Epidermis-20, Intestine-9, Intestine-12, Muscle-6, and Muscle-16 populations. Each published population was matched to the updated subcluster containing the largest fraction of its cells. Early regenerative enrichment was assessed by calculating, for every updated subcluster represented in the unirradiated dataset, the fraction of cells originating from days 0, 1, 2, and 4. Candidate TRACS were defined by an early-time-point fraction exceeding the mean across subclusters by more than two standard deviations. Refined TRACS correspondence required distinct matched subclusters within epidermis, intestine, or muscle that met this criterion. Occupancy was calculated for each treatment and time point as the percentage of captured cells within the corresponding tissue lineage. Published TRACS occupancy used the original tissue annotations, whereas refined TRACS occupancy used the updated annotations. Subcluster abundance was also calculated relative to all captured cells. The composition of updated Muscle_9 was additionally examined by separating cells originating from published Muscle-6, Muscle-16, or other subclusters.

Activin-1 (SMED30031091), Activin-2 (SMED30035469), Activin-3 (SMED30009429), and Follistatin (SMED30035357) expression was quantified using the RNA assay. Raw counts were normalized to 10,000 counts per cell and natural-log transformed with a pseudocount of one using Seurat’s LogNormalize method. Expression values were explicitly extracted from the RNA assay data layer. For each treatment and time point, we calculated the percentage of cells with detected expression, mean log-normalized expression across all cells, and mean log-normalized expression among positive cells. Positive cells were defined by expression values greater than zero, corresponding to nonzero raw RNA counts. Summaries were calculated across all captured cells and separately within muscle and intestine.

Positive markers for updated subclusters were identified within each tissue lineage and against all other subclusters in the global object using Seurat’s FindAllMarkers function with the Wilcoxon rank-sum test. Analyses used the RNA assay data layer, applying LogNormalize normalization with a scale factor of 10,000 when this layer was absent. Parameters were min.pct = 0.10, logfc.threshold = 0.25, only.pos = TRUE, and return.thresh = 0.05. Marker tables were annotated using SMED gene annotations. Analysis code accompanies the manuscript.

### Statistical Analyses

For morphological phenotype significance, individual worm scores (0-4) were compared to the control group (Un-irradiated or *Unc-22* RNAi). Statistical significance for phenotype differences was determined using a Mann-Whitney test in GraphPad Prism 10 (v10.2.3) with correction for multiple comparisons. For CNS and pharynx regeneration, the areas of CNS or pharynx tissue in each individual worm were measured using FIJI ImageJ image analysis tools, then were compared to those of the unc-22-RNAi-treated control. In all cases, each variable’s individuals and sample sizes were used for independent comparisons to the control, with heads, trunk, and tails analyzed separately. Statistical significance was determined using a Mann-Whitney test in GraphPad Prism 10 (v10.2.3) with correction for multiple comparisons. For quantitation of H3P density, % piwi+ cells, and % piwi+ cells H3P+, statistical significance was determine using a Student T-test in GraphPad Prism 10 (v10.2.3) with corrections for multiple comparisons.

### Cisplatin treatment

A 10mM cisplatin stock solution was created by dissolving Enzo cisplatin powder in milli-Q water. Planarians were prepared for treatment in 6-well plates containing 1x Montjuic planarian water. A volume of stock solution was added to each well to create final cisplatin concentrations. Cisplatin-treated animals were soaked in treatment for 24 hours before being relocated to new plates in 1x Montjuic + gentamicin (50ug/ml).

### Irradiation of planarians

Planarians were dosed with gamma irradiation in a Cesium-137 self-shielded low dose-rate blood irradiator. Following irradiation planarians were maintained in petri dishes in 1x Montjuic + gentamicin (50ug/ml).

### Re-injury after irradiation assay

Irradiated head and tail regenerates 14dpa were generated using the previously described amputation method, cut 7 days after 750R and 1250R doses. To reintroduce injury, head regenerates were cut at the posterior wound site, while tails were cut at the anterior wound site. For regenerates with no blastema formation, the healed wound site was cut. For regenerates with any degree of blastema formation, the blastema was amputated.

## Notes

### Competing Interest Statement

The authors have declared no competing interest.

### Summary of Updates

- additional analyses of TRACS trajectories in WT vs. irradiated worms - addition of experiments visualizing activin, follistatin, and notum expression in irradiated worms after amputation - addition of re-cut experiments demonstrating improved regeneration if worms are re-cut after failed regeneration - addition of experiments demonstrating that cisplatin treatment also inhibits regeneration and can be rescued by activin-2 depletion - additional phenotypic characterization of activin-2 RNAi animals.

