## Supplementary Figures for "Tissue-derived Activin signaling restrains regeneration after genotoxic stress in *Schmidtea mediterranea*"

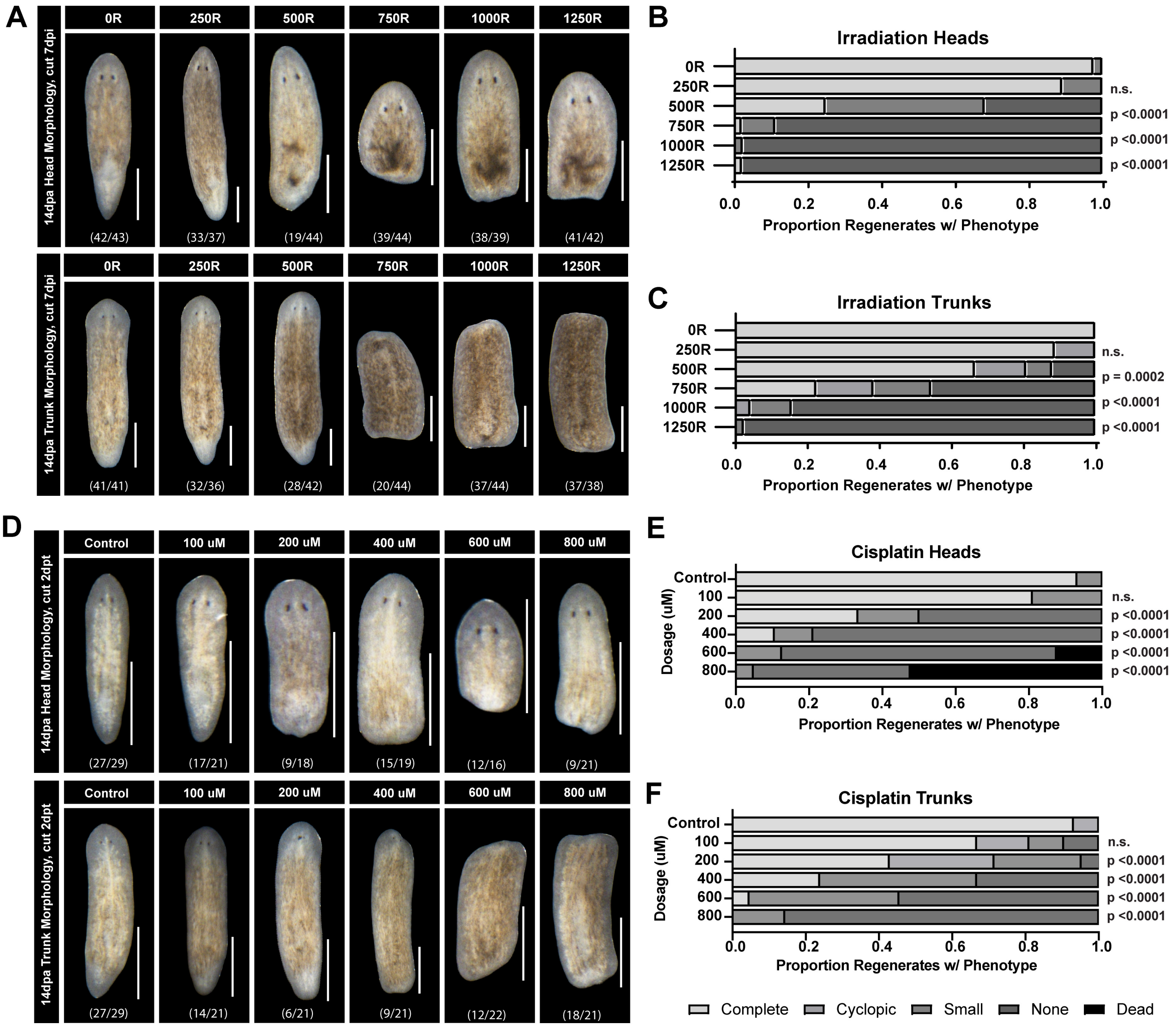


**Figure S1. Inhibition of regeneration in heads and trunks after low dose genotoxic stress**
(A) Morphologies of heads (top) and trunk (bottom) regenerates 14dpa when cut 7 days post irradiation. Quantitation of head (B) and trunk (C) phenotypes at irradiation doses ranging from 0-1250R. (D) Morphologies of heads (top) and trunk (bottom) regenerates 14dpa when cut 2 days post cisplatin soak. Quantitation of head (E) and trunk (F) phenotypes at cisplatin soak doses ranging from 0-800uM. P-values are Mann-Whitney test with correction for multiple comparisons. Scale = 1mm.

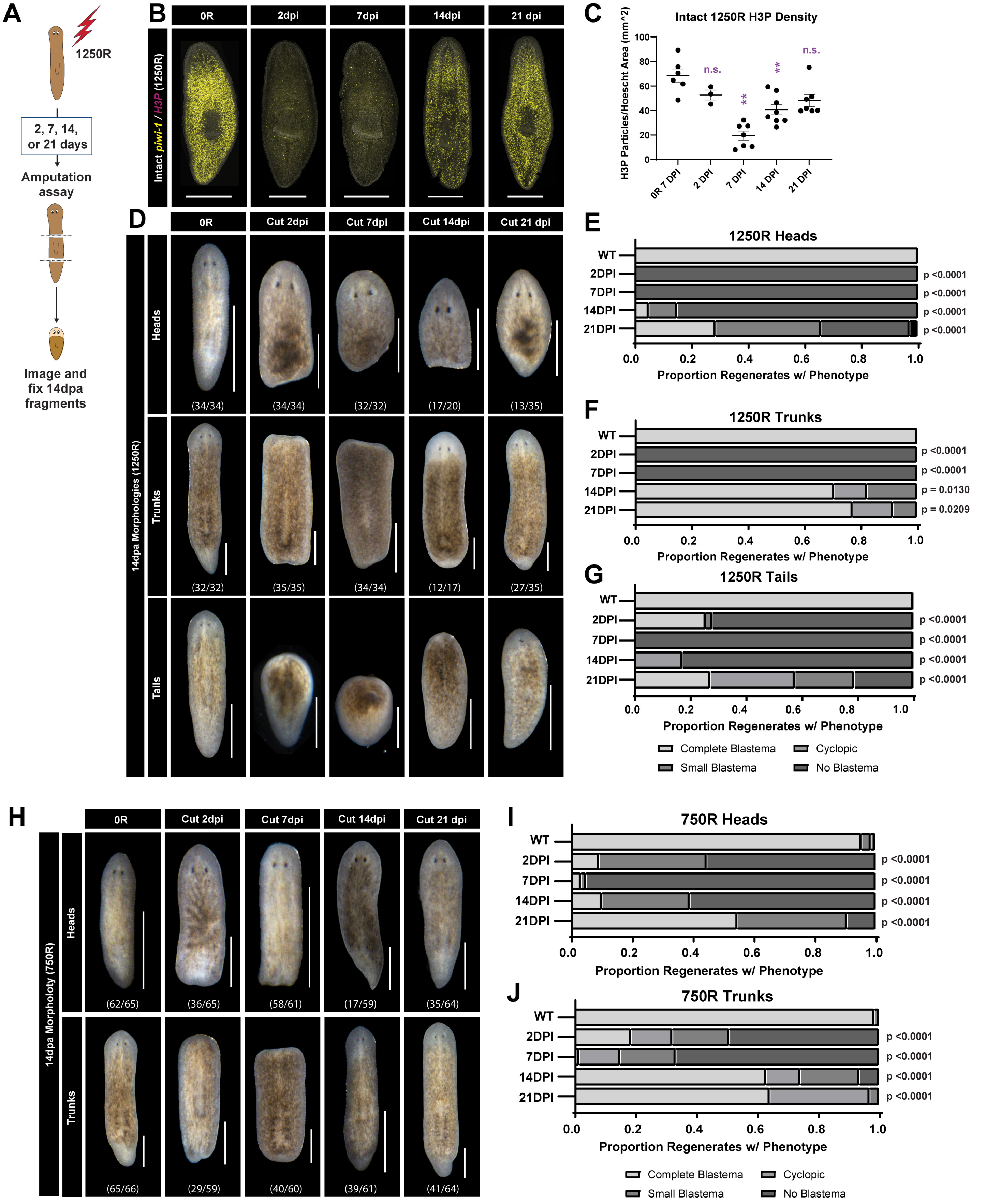


**Figure S2. Inhibition of regeneration is transient after both 750 and 1250R**

(A) Schematic of irradiation time course experiment (1250R) (B) Whole mount *piwi-1* in situ hybridization merged with H3P antibody staining at 0, 2, 7, 14, and 21dpi (C) Scatter plot of H3P density quantitation with SEM bars at 0, 2, 7, 14, and 21dpi (D) Morphologies of 1250R head (top), trunk (middle), and tail (bottom) regenerates 14dpa. Quantitation of 1250R head (E), trunk (F), and tail (G) regeneration phenotypes at 0, 2, 7, 14, and 21dpi (H) Morphologies of 750R head (top) and trunk (bottom) regenerates at 14dpa. Quantitation of 750R head (I) and trunk (J) regeneration phenotypes at 0, 2, 7, 14, and 21dpi. P-values are Mann-Whitney test with correction for multiple comparisons. Scale = 1mm.

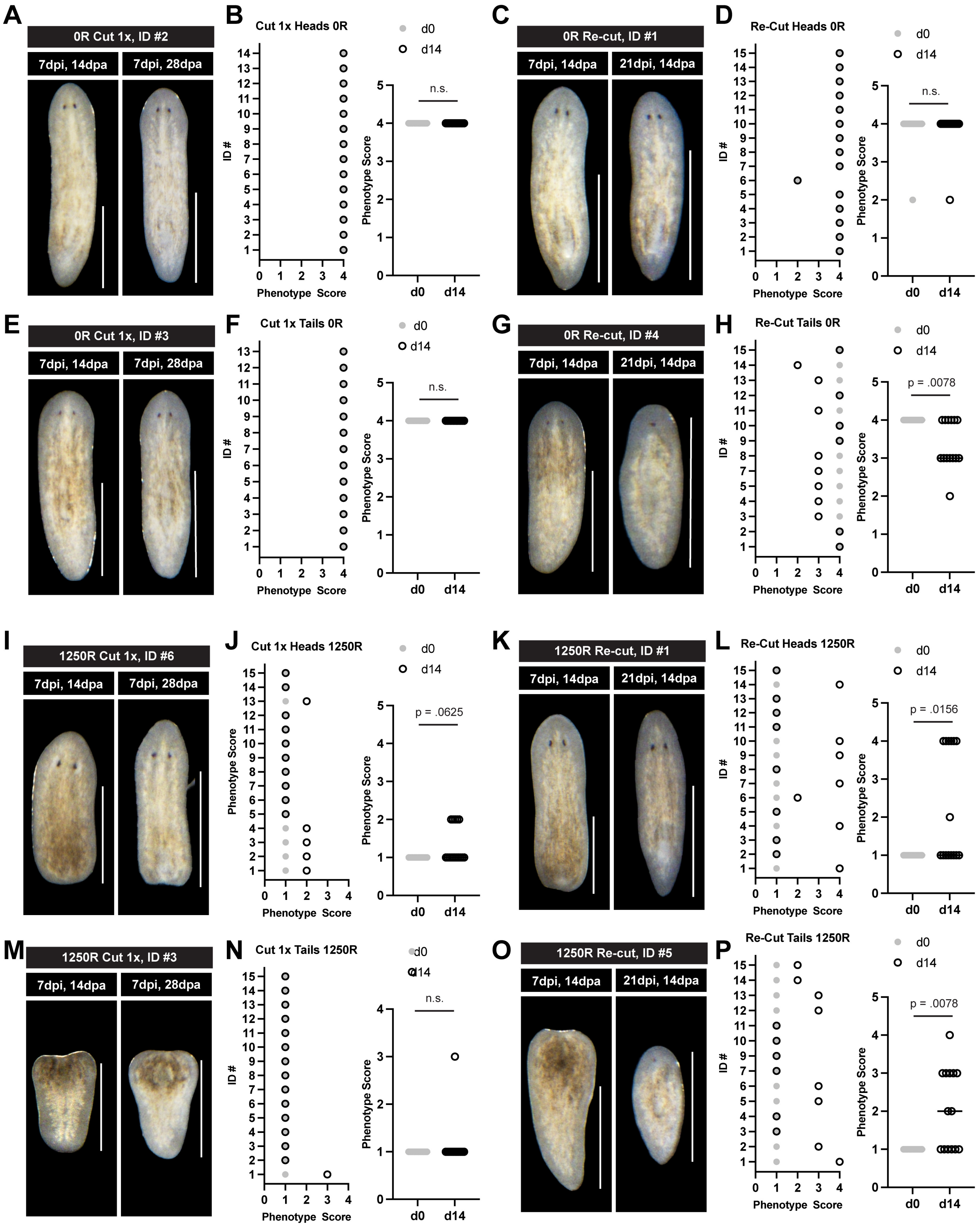

A
0R Cut 1x, ID #2
7dpi, 14dpa
7dpi, 28dpa
B
Cut 1x Heads 0R
ID #
Phenotype Score
d0
d14
n.s.
C
0R Re-cut, ID #1
7dpi, 14dpa
21dpi, 14dpa
D
Re-Cut Heads 0R
n.s.
E
0R Cut 1x, ID #3
7dpi, 14dpa
7dpi, 28dpa
F
Cut 1x Tails 0R
n.s.
G
0R Re-cut, ID #4
7dpi, 14dpa
21dpi, 14dpa
H
Re-Cut Tails 0R
p = .0078
I
1250R Cut 1x, ID #6
7dpi, 14dpa
7dpi, 28dpa
J
Cut 1x Heads 1250R
p = .0625
K
1250R Re-cut, ID #1
7dpi, 14dpa
21dpi, 14dpa
L
Re-Cut Heads 1250R
p = .0156
M
1250R Cut 1x, ID #3
7dpi, 14dpa
7dpi, 28dpa
N
Cut 1x Tails 1250R
n.s.
O
1250R Re-cut, ID #5
7dpi, 14dpa
21dpi, 14dpa
P
Re-Cut Tails 1250R
p = .0078

**Figure S3. Re-injury phenotypes in animals treated with 0R or 1250R prior to amputation. (A)** Head regeneration morphologies of unirradiated head #2 cut 7dpi, imaged at 14dpa (left), and imaged again at 28dpa (right). (B) Scatter plot of individual unirradiated head phenotype scores cut 7dpi, quantified at 14dpa (d0) and 28dpa (d14) (left), Head phenotype scores cut 7dpi, quantified at d0 vs d14 (right). (C) Head regeneration morphologies of unirradiated head #1 cut 7dpi, imaged at 14dpa (left), then cut again at 21dpi, imaged 14dpa (right). (D) Scatter plot of individual head phenotype scores cut 7dpi, quantified at 14dpa (d0) and cut again at 21 dpi, quantified 14dpa (d14) (left), Head phenotype scores at d0 vs d14. (E) Tail regeneration morphologies of unirradiated tail #3 cut 7dpi, imaged at 14dpa (left), and imaged again at 28dpa (right) (F) Scatter plot of individual tail phenotype scores cut 7dpi, quantified at 14dpa (d0) and 28dpa (d14) (left), Tail phenotype scores cut 7dpi, quantified at d0 vs d14 (right). (G) Tail regeneration morphologies of unirradiated tail #4 cut 7dpi, imaged at 14dpa (left), then cut again at 21dpi, imaged 14dpa (right). (H) Scatter plot of individual tail phenotype scores cut 7dpi, quantified at 14dpa (d0) and cut again at 21 dpi, quantified 14dpa (d14) (left), Tail phenotype scores at d0 vs d14. (I) Head regeneration morphologies of 1250R head #6 cut 7dpi, imaged at 14dpa (left), and imaged again at 28dpa (right). (J) Scatter plot of 1250R head phenotype scores cut 7dpi, quantified at 14dpa (d0) and 28dpa (d14) (left), Head phenotype scores cut 7dpi, quantified at d0 vs d14 (right). (K) Head regeneration morphologies of 1250R head #1 cut 7dpi, imaged at 14dpa (left), then cut again at 21dpi, imaged 14dpa (right) (L) Scatter plot of individual head phenotype scores cut 7dpi, quantified at 14dpa (d0) and cut again at 21 dpi, quantified 14dpa (d14) (left), Head phenotype scores at d0 vs d14. (M) Tail regeneration morphologies of 1250R tail #3 cut 7dpi, imaged at 14dpa (left), and imaged again at 28dpa (right) (N) Scatter plot of individual tail phenotype scores cut 7dpi, quantified at 14dpa (d0) and 28dpa (d14) (left), Tail phenotype scores cut 7dpi, quantified at d0 vs d14 (right). (O) Tail regeneration morphologies of 1250R tail #5 cut 7dpi, imaged at 14dpa (left), then cut again at 21dpi, imaged 14dpa (right). (P) Scatter plot of individual tail phenotype scores cut 7dpi, quantified at 14dpa (d0) and cut again at 21 dpi, quantified 14dpa (d14) (left), Tail phenotype scores at d0 vs d14. P values are Mann-Whitney test with correction for multiple comparisons. Scale = 1mm.

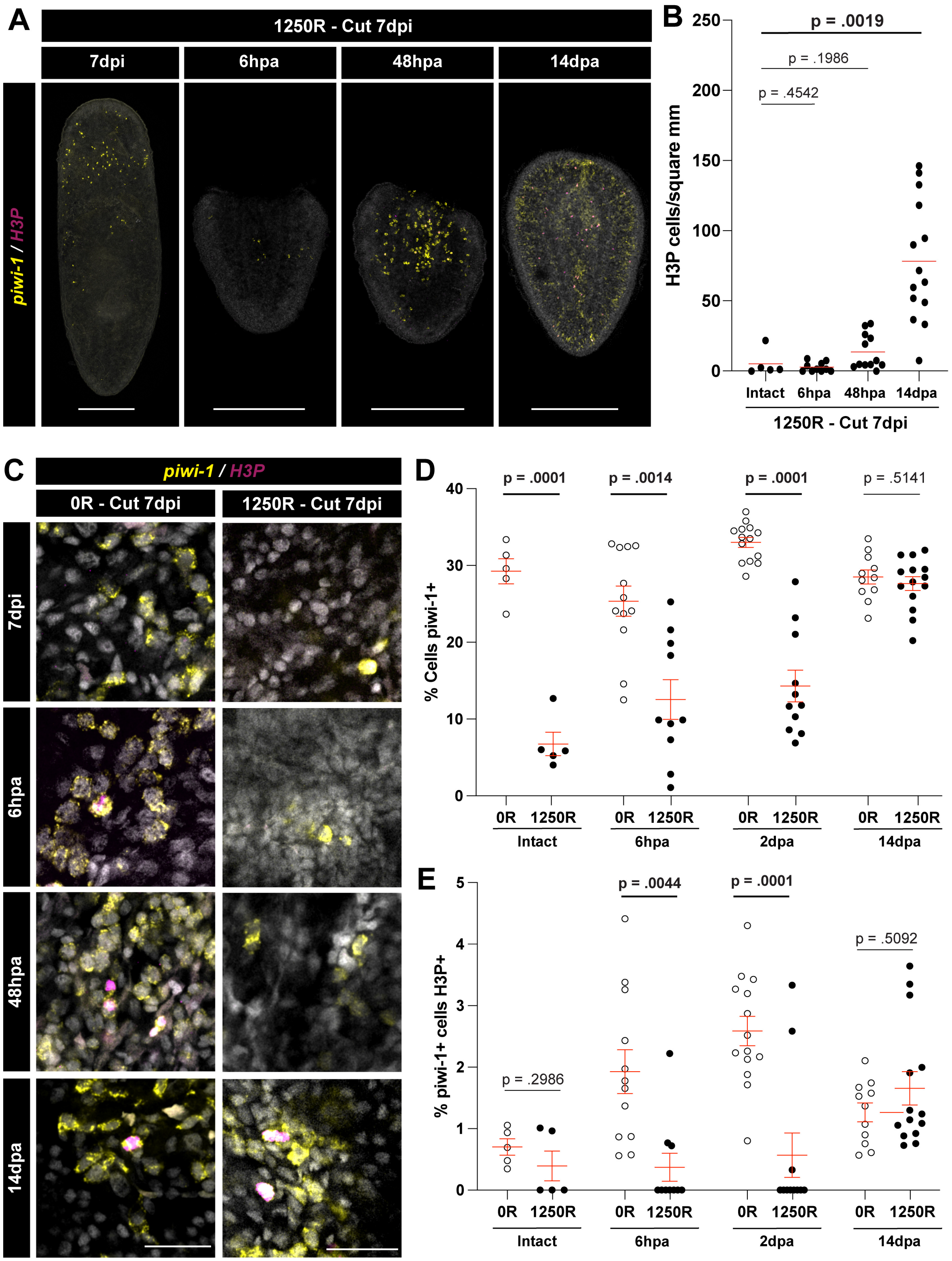


**Figure S4. Stem cell numbers and proliferation in regenerates after 1250R exposure**

(A) Whole mount max projections of intact and regenerating animals 6hpa, 48hpa, or 14dpa stained with *piwi-1* and H3P if cut 7dpi with 1250R (B) Quantitation of H3P density in the same worms shown in A (C) Confocal slices of ROIs from animals in A showing *piwi-1* and H3P staining. Quantitation of percentage of cells *piwi-1*$^{+}$ (D) and percentage of *piwi-1*$^{+}$ cells also H3P$^{+}$ (E) in animals from C. P-values are Mann-Whitney test with correction for multiple comparisons. Scale = 1mm (A) or 25uM (C).

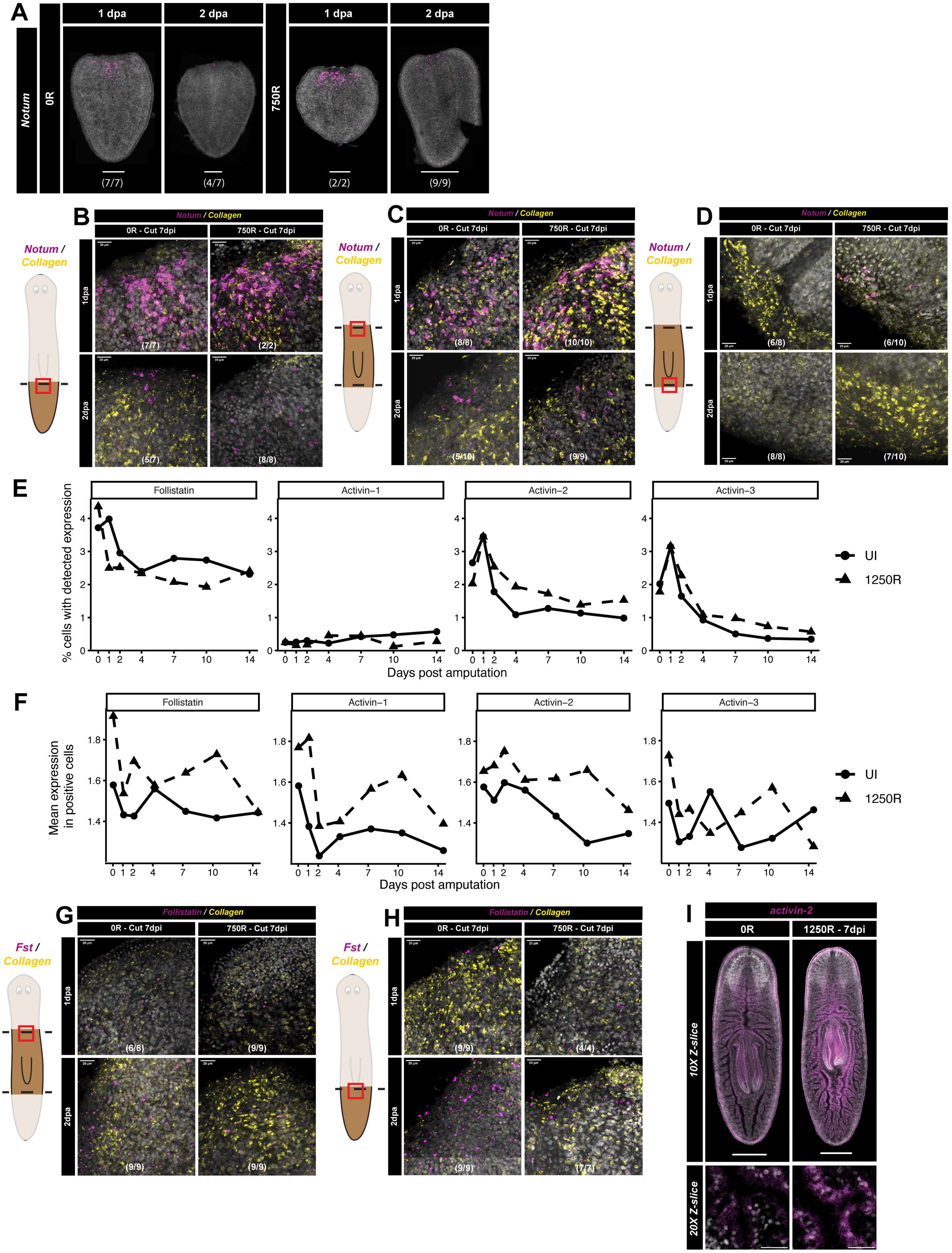

A
Notum
0R
750R
1 dpa
2 dpa
(7/7)
(4/7)
(2/2)
(9/9)
B
Notum / Collagen
0R - Cut 7dpi
750R - Cut 7dpi
1dpa
2dpa
(7/7)
(2/2)
(5/7)
(8/8)
C
(8/8)
(10/10)
(5/10)
(9/9)
D
(6/8)
(6/10)
(8/8)
(7/10)
E
Follistatin
Activin−1
Activin−2
Activin−3
% cells with detected expression
Days post amputation
UI
1250R
F
Mean expression in positive cells
G
Follistatin / Collagen
Fst / Collagen
(6/6)
(9/9)
(9/9)
(9/9)
H
(9/9)
(4/4)
(9/9)
(7/7)
I
activin-2
0R
1250R - 7dpi
10X Z-slice
20X Z-slice

**Figure S5. Wound-induced expression of *notum, follistatin,* and *activin* ligands in regenerates from unirradiated and low-dose irradiated worms.**
(A) Whole mount max projections of regenerating tails cut at 7dpi fixed at 1 or 2dpa stained with *notum* via FISH. 20x max projections of regenerating tails (B), anterior facing trunk wounds (C), or posterior facing trunks wounds (D) fixed at 1 or 2dpa, visualizing *collagen* and *notum* via double FISH. (E) Percent total cells expressing *follistatin* and *activin* ligands in unirradiated or 1250R treated regeneration time courses. (F) Normalized expression of *follistatin* and *activin* ligands in positive cells captured from unirradiated or 1250R treated regeneration time courses. 20x max projections of regenerating anterior trunk wounds (G) or tails (H) fixed at 1 or 2dpa, visualizing collagen and follistatin via double FISH. (I) Whole mount max projections of Intact worms fixed at 7dpi stained with *activin-2* via FISH.

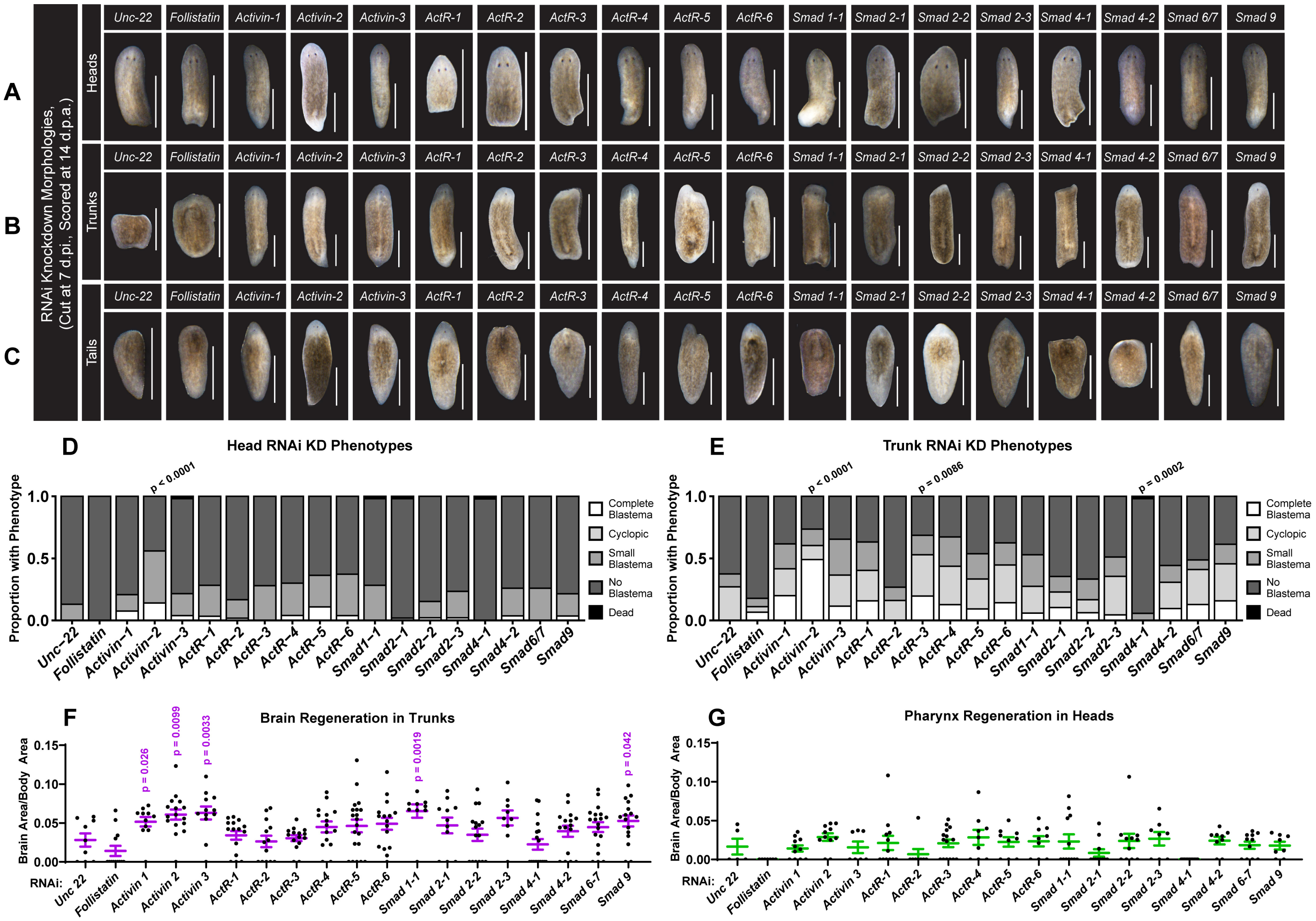

**Figure S6. An RNAi screen of activin homologs identifies regulators of regenerative capacity after low dose irradiation.**

Morphologies of RNAi-treated head (A), trunk (B) and tail (C) regenerates 14dpa, cut 7 days post 750R treatment. Phenotype quantitation of head (D) and trunk (E) RNAi-treated fragments 14dpa, cut 7 days post 750R treatment. Quantitation of brain size in regenerating trunks (F) and pharynx size in regenerating heads (G) cut at 7dpi quantified at 14dpa. P-values are Mann-Whitney test with correction for multiple comparisons. Scale = 1mm.

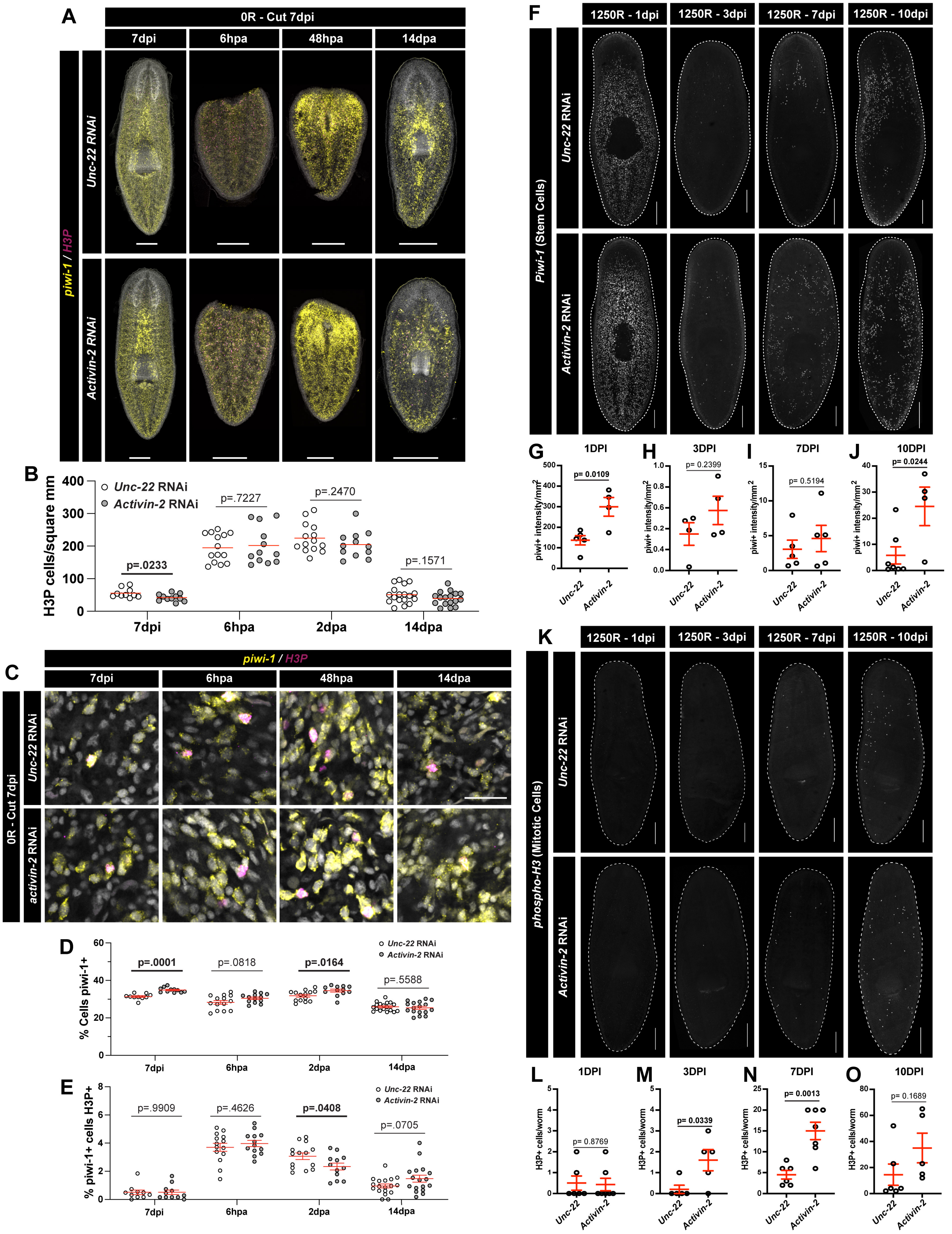

A
0R - Cut 7dpi
7dpi
6hpa
48hpa
14dpa
piwi-1 / H3P
Unc-22 RNAi
Activin-2 RNAi
B
H3P cells/square mm
Unc-22 RNAi
Activin-2 RNAi
p=.0233
p=.7227
p=.2470
p=.1571
7dpi
6hpa
2dpa
14dpa
C
piwi-1 / H3P
0R - Cut 7dpi
Unc-22 RNAi
activin-2 RNAi
D
% Cells piwi-1+
p=.0001
p=.0818
p=.0164
p=.5588
E
% piwi-1+ cells H3P+
p=.9909
p=.4626
p=.0408
p=.0705
F
1250R - 1dpi
1250R - 3dpi
1250R - 7dpi
1250R - 10dpi
Piwi-1 (Stem Cells)
G
1DPI
p= 0.0109
H
3DPI
p= 0.2399
I
7DPI
p= 0.5194
J
10DPI
p= 0.0244
piwi+ intensity/mm2
Unc-22
Activin-2
K
phospho-H3 (Mitotic Cells)
L
1DPI
p= 0.8769
M
3DPI
p= 0.0339
N
7DPI
p= 0.0013
O
10DPI
p= 0.1689
H3P+ cells/worm

**Figure S7. *Activin-2*-dependent stem cell proliferation and regeneration responsiveness in worms treated with 1250R.**

(A) Whole mount max projections of intact and regenerating RNAi-treated animals 6hpa, 48hpa, or 14dpa when cut 7dpi treatment with 1250R, stained with *piwi-1* and H3P (B) Quantitation of H3P density in the same worms shown in A (C) Confocal slices of ROIs from animals in A showing *piwi-1* and H3P staining. Quantitation of percentage of cells *piwi-1*$^{+}$ (D) and percentage of *piwi-1*$^{+}$ cells also H3P$^{+}$ (E) in ROIs from C. (F) Whole mount max projections of *piwi-1* expression by FISH in intact RNAi-treated animals 1-14 days post 1250R treatment. Quantitation of piwi-1 intensity per area at 1dpi (G), 3dpi (H), 7dpi (I), and 14dpi (J). (K) Whole mount max projections of Phospho-H3 by immunohistochemistry in intact RNAi-treated animals 1-14 days post 1250R treatment. Quantitation of H3P density at 1dpi (L), 3dpi (M), 7dpi (N), and 14dpi (O). P-values are Mann-Whitney test with correction for multiple comparisons. Scale = 1mm (A), 500uM (F, K) or 25uM (C).

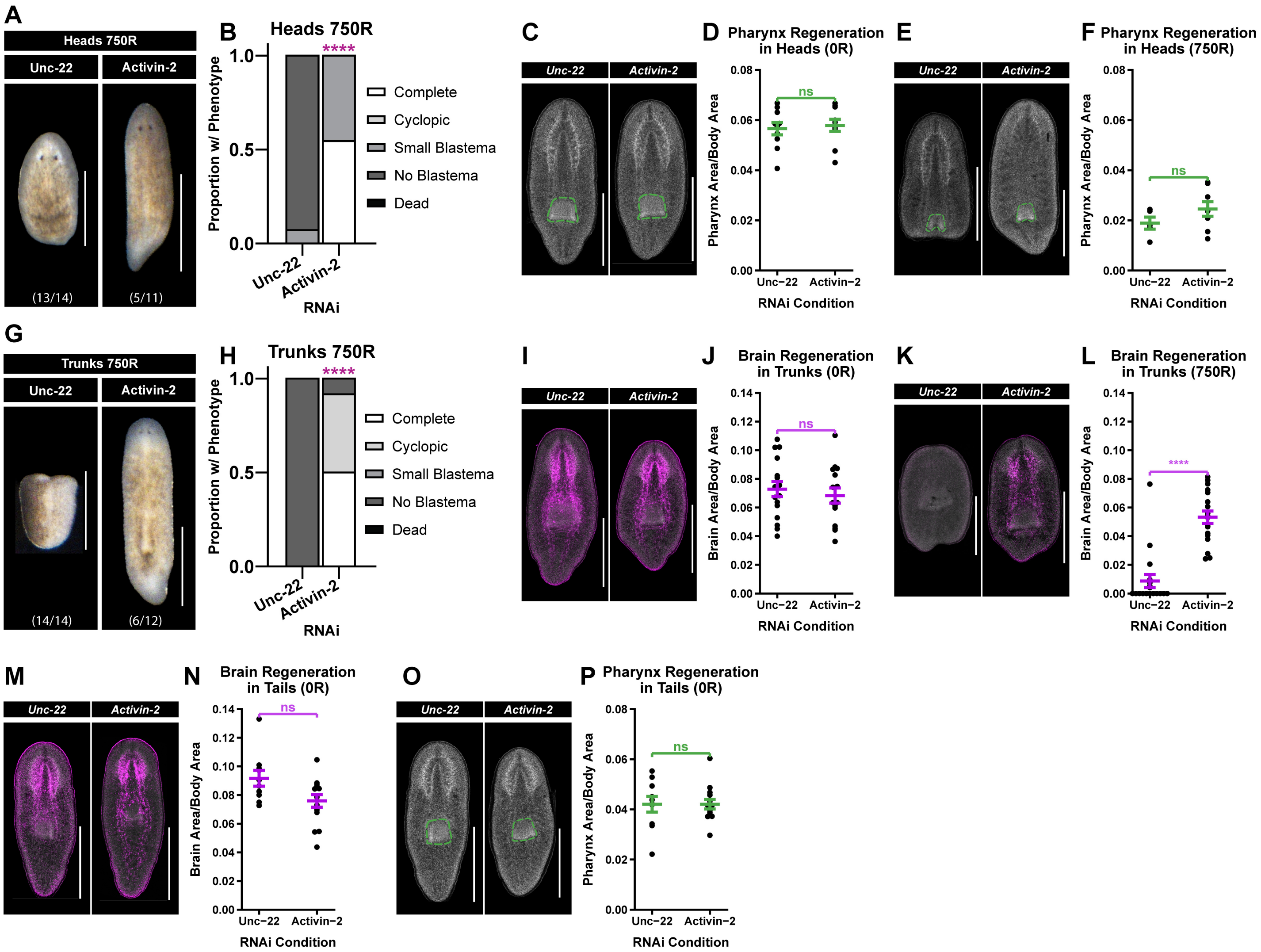


**Figure S8. Additional characterization of *activin-2*-dependent regeneration phenotypes**

(A) Morphologies of 750R *unc-22* RNAi controls vs *activin-2* RNAi-treated head regenerates 14dpa. (B) Quantitation of regeneration phenotypes from animals in A. Whole mount max projections of Hoechst staining, visualizing pharynx size (green) in RNAi-treated regenerating heads 14dpa, cut 7 days after 0R (C) or 750R (E). Quantitation of pharynx size in *unc-22* RNAi controls vs *activin-2* RNAi-treated animals 14dpa, cut 7 days after 0R (D) or 750R (F). (G) Morphologies of 750R treated RNAi trunk regenerates 14dpa. Whole mount max projections of *prohormone convertase-2* (PC2) staining visualizing brain size (magenta) in RNAi-treated regenerating trunks 14dpa, cut 7 days after 0R (I) or 750R (K). Quantitation of brain size in *unc-22* RNAi controls vs *activin-2* RNAi-treated animals 14dpa, cut 7 days after 0R (J) or 750R (L). (M) Max projections of PC2 staining visualizing brain size in RNAi-treated regenerating tails 14dpa, cut 7 days after 0R. (N) Quantitation of brain size after RNAi treatment in regenerating tails 14dpa, cut 7 days after 0R. (O) Max projections of Hoechst staining, visualizing pharynx size in RNAi-treated regenerating tails 14dpa, cut 7 days after 0R. P-values are Mann-Whitney test with correction for multiple comparisons. Scale = 1mm.

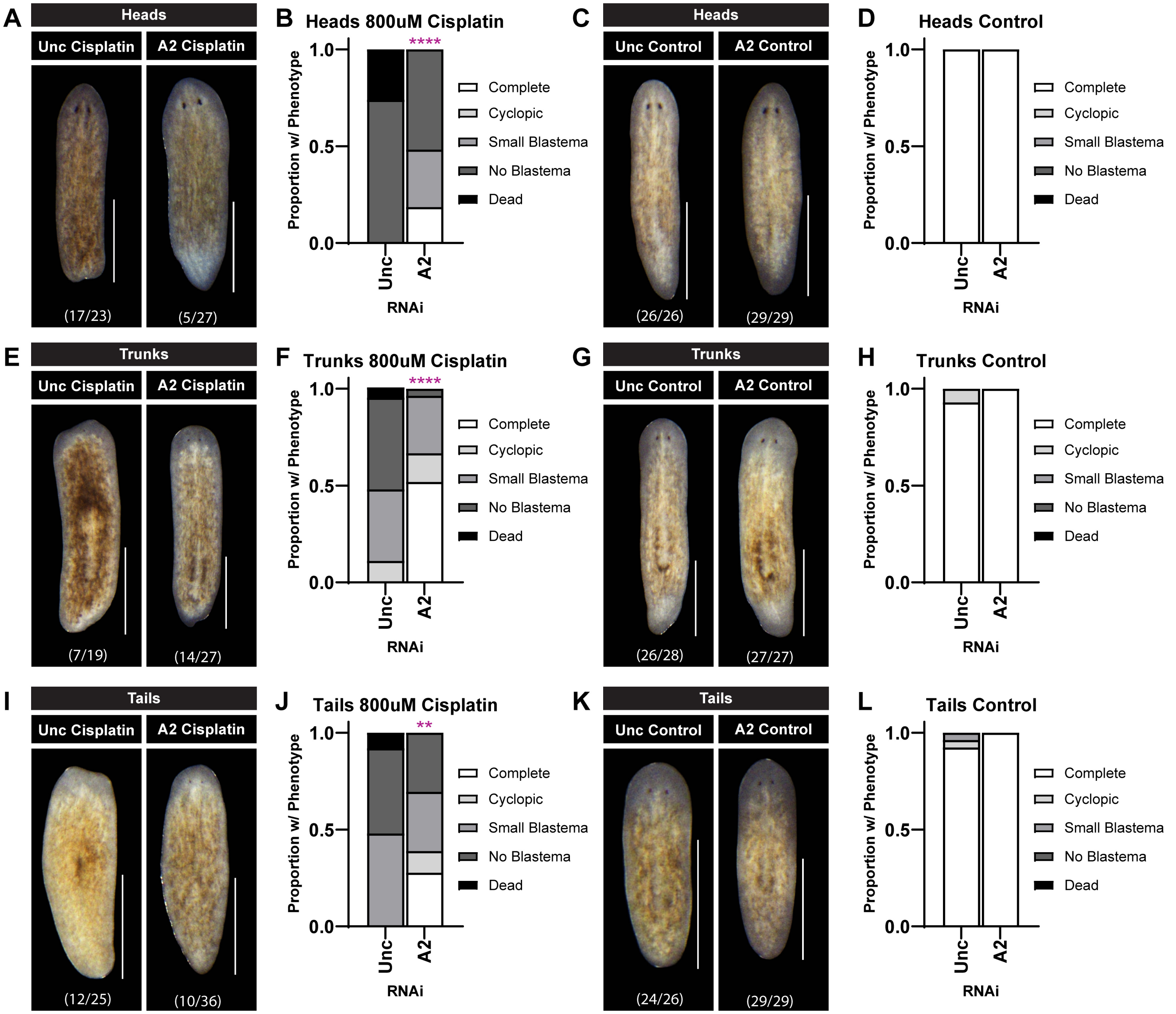


**Figure S9. Activin-2 depletion rescues regeneration following cisplatin treatment**
Morphologies of RNAi-treated regenerating heads (A), trunks (E), and tails (I) 14dpa, cut 2 days after cisplatin treatment. Quantitation of regeneration phenotypes in cisplatin-treated heads (B), trunks (F), and tails (J) comparing Unc-22 (control) vs Activin-2 RNAi treatment. Morphologies of RNAi-treated regenerating heads (C), trunks (G), and tails (K) 14dpa. Quantitation of regeneration phenotypes in water control heads (D), trunks (H), and tails (L) comparing Unc-22 (control) vs Activin-2 RNAi treatment. P-values are Mann-Whitney test with correction for multiple comparisons. Scale = 1mm.

### SUPPLEMENTARY TABLES

**Table 1. List of genes in the study and corresponding SMEDIDs from planosphere database**
